# Hydrogen-dependent dissimilatory nitrate reduction to ammonium enables growth of *Campylobacterota* isolates

**DOI:** 10.1101/2024.09.06.611579

**Authors:** Hokwan Heo, Thanh Nguyen-Dinh, Man-Young Jung, Chris Greening, Sukhwan Yoon

## Abstract

Dissimilatory nitrate reduction to ammonium (DNRA) is a key process in global nitrogen cycling, supporting the energy conservation of diverse microbes. For a long time, DNRA has been thought to primarily depend on organic electron donors, and thus to be governed by carbon-to-nitrogen (C:N) ratios. However, recent studies suggest that inorganic electron donors, such as sulfur compounds and iron, may also facilitate DNRA. The coupling of DNRA with molecular hydrogen (H_2_) oxidation is theoretically feasible, but largely unexplored microbial process. Here, we report the isolation of two *Campylobacterota* strains, *Aliarcobacter butzleri* hDNRA1 and *Sulfurospirillum* sp. hDNRA2, that grow by using H_2_ as an electron donor for DNRA. In both batch and continuous cultures, DNRA *sensu stricto*, i.e., NO_2−_-to-NH_4_ ^+^ reduction, depended on the presence of H_2_ and was stoichiometric with H_2_ oxidation. The electrons for NO_2_ ^−^ reduction were clearly derived from H_2_, and hydrogenotrophic DNRA was largely unaffected by the ratio of either carbon or electron donor to NO_3_ ^−^/NO_2_ ^−^. Genomic and transcriptomic analyses indicate that group 1b [NiFe]-hydrogenase and cytochrome *c*_552_ nitrite reductase are the key enzymes catalyzing hydrogenotrophic DNRA. These findings reveal novel physiological mechanisms enabling anaerobic bacterial growth, challenge the traditional C:N ratio paradigm, and uncover new biogeochemical processes and mediators controlling the global nitrogen and hydrogen cycles.

## INTRODUCTION

Nitrogen is an essential element for the sustenance and growth of all life forms (1). While nitrogen is an indispensable nutrient, the excessive release of anthropogenic reactive nitrogen into the environment poses significant threats to global sustainability by exacerbating nutrient pollution and climate change (2–4). The transformation of one nitrogen form to another, largely mediated by diverse microorganisms, dictates nitrogen availability to primary producers, both when nitrogen abundance is beneficial (e.g., agricultural soils) and detrimental (e.g., eutrophic water bodies) (5, 6). Microbial nitrogen-cycling reactions, including nitrification (aerobic NH_4_ ^+^ oxidation to NO_3_ ^−^ via NO_2_ ^−^), denitrification (anaerobic NO_3_ ^−^ reduction to NO, N_2_O, and N_2_), and anammox (anaerobic NH_4_ ^+^ oxidation to N_2_), have been widely studied in this context, and also due to their contributions to the emission of the potent greenhouse gas N_2_O into the atmosphere (7–9).

Another biological nitrogen-cycling reaction, dissimilatory nitrate/nitrite reduction to ammonium (DNRA), also known as respiratory nitrate ammonification, remains relatively understudied despite its potential environmental significance. As with denitrification, DNRA is an anaerobic bacterial respiratory pathway that reduces NO_3_ ^−^ via NO_2_ ^−^ for energy conservation (10). What distinguishes DNRA from denitrification is the reaction step that reduces NO_2_ ^−^ to NH_4_ ^+^, typically catalyzed by cytochrome *c*_552_ nitrite reductases (NrfA) (11, 12). From an environmental perspective, DNRA’s competition with denitrification for NO_3_ ^−^/NO_2_ ^−^ is particularly significant, as DNRA results in nitrogen retention, thereby preventing or alleviating nitrogen loss caused by denitrification (5, 11). Previous studies have tried to explain this ecological competition as being primarily influenced by the C:N ratio, i.e., the ratio of organic electron donors over nitrogenous electron acceptors (13, 14). For decades, DNRA has been hypothesized to be favored in environments with high C:N ratios due to its higher per-electron-acceptor energy yield, and numerous field and laboratory observations have supported this hypothesis (13–17).

Recent studies have shown that DNRA can also be coupled with the oxidation of inorganic electron donors, i.e., chemolithotrophic DNRA, bypassing the C:N ratio hypothesis and thereby complicating the ecological implications of DNRA. Reduced forms of sulfur, i.e., elemental sulfur (S^0^) and sulfide (H_2_S/HS^−^/S^2−^), have been suggested as potential electron donors for chemolithotrophic DNRA, explaining observations of elevated DNRA activity in sulfur-rich, highly reduced aquatic sediments (18–20). The coupling of DNRA with ferrous iron (Fe^2+^) oxidation has also been proposed based on the studies with cable bacteria-amended sediments and anammox enrichments (21, 22). Chemolithotrophic DNRA has been observed in several axenic cultures as well, e.g., *Desulfurivibrio alkaliphilus* using sulfide as the electron donor. However, the current understanding is too limited to draw definite conclusions about its biogeochemical significance (23, 24). Particularly puzzling is the paucity of investigations on DNRA coupling with the oxidation of molecular hydrogen (H_2_), given that this reaction is highly exergonic (equations 1 and 2), H_2_ is a ubiquitous electron donor, and respiratory hydrogenases are widely distributed across microbiomes in diverse ecosystems (25–27).

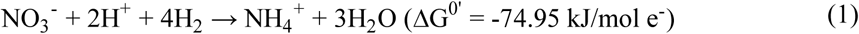

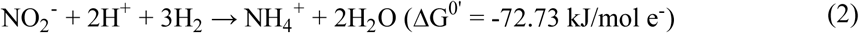

Only one previous study has provided strong evidence of hydrogenotrophic DNRA, demonstrating that *Desulfovibrio desulfuricans* can grow via DNRA in the presence of H_2_ (28). However, the physiological dependence of DNRA on H_2_ oxidation was not examined, and the study, which preceded the genomic era, lacked a genomic or enzymatic basis for the observed phenotype. Here, as part of our efforts to isolate DNRA-catalyzing microorganisms from activated sludge samples (16), we inadvertently isolated two bacteria capable of conserving energy through the coupling of H_2_ oxidation to NO_3_ ^−^-to-NH_4_ ^+^ reduction (hereafter referred to as hydrogenotrophic DNRA). By integrating physiological, genomic, and transcriptomic analyses, we confirmed the dependence of DNRA on H_2_ and identified the key genes involved in this metabolism. These findings provide novel insights into DNRA ecophysiology, expand our understanding of microbial metabolic diversity, and challenge the prevailing paradigm that DNRA activity is primarily governed by the C:N ratio and/or redox potential of the environment.

## RESULTS

### New *Campylobacterota* isolates mediate hydrogenotrophic DNRA

We isolated two bacteria that mediated NO_3_ ^−^/NO_2_ ^−^ reduction to NH_4_ ^+^ only in the presence of H_2_. Genomic analyses showed these strains were phylogenetically affiliated with the genera *Aliarcobacter* and *Sulfurospirillum*, both belonging to the phylum *Campylobacterota* (Fig. 1A). Their closest type strains, based on both 16S rRNA gene and whole-genome sequences, were identified as *Aliarcobacter butzleri* RM4018 (99.9% 16S rRNA gene sequence identity; digital DNA-to-DNA hybridization score of 76.1% using the formula *d_4_*) and *Sulfurospirillum cavolei* NBRC 109482 (99.5% and 67.0%, respectively) (29). Thus, the two new isolates are hereafter referred to as *Aliarcobacter butzleri* hDNRA1 and *Sulfurospirillum* sp. hDNRA2. Both isolates exhibited a curved rod-shaped morphology with approximate dimensions of 1.5-2.5 µm × 0.3-0.4 µm (Fig. 1B). Although their morphologies were very similar, only *A*. *butzleri* hDNRA1 possessed a polar flagellum-like appendage.

**Fig. 1.**
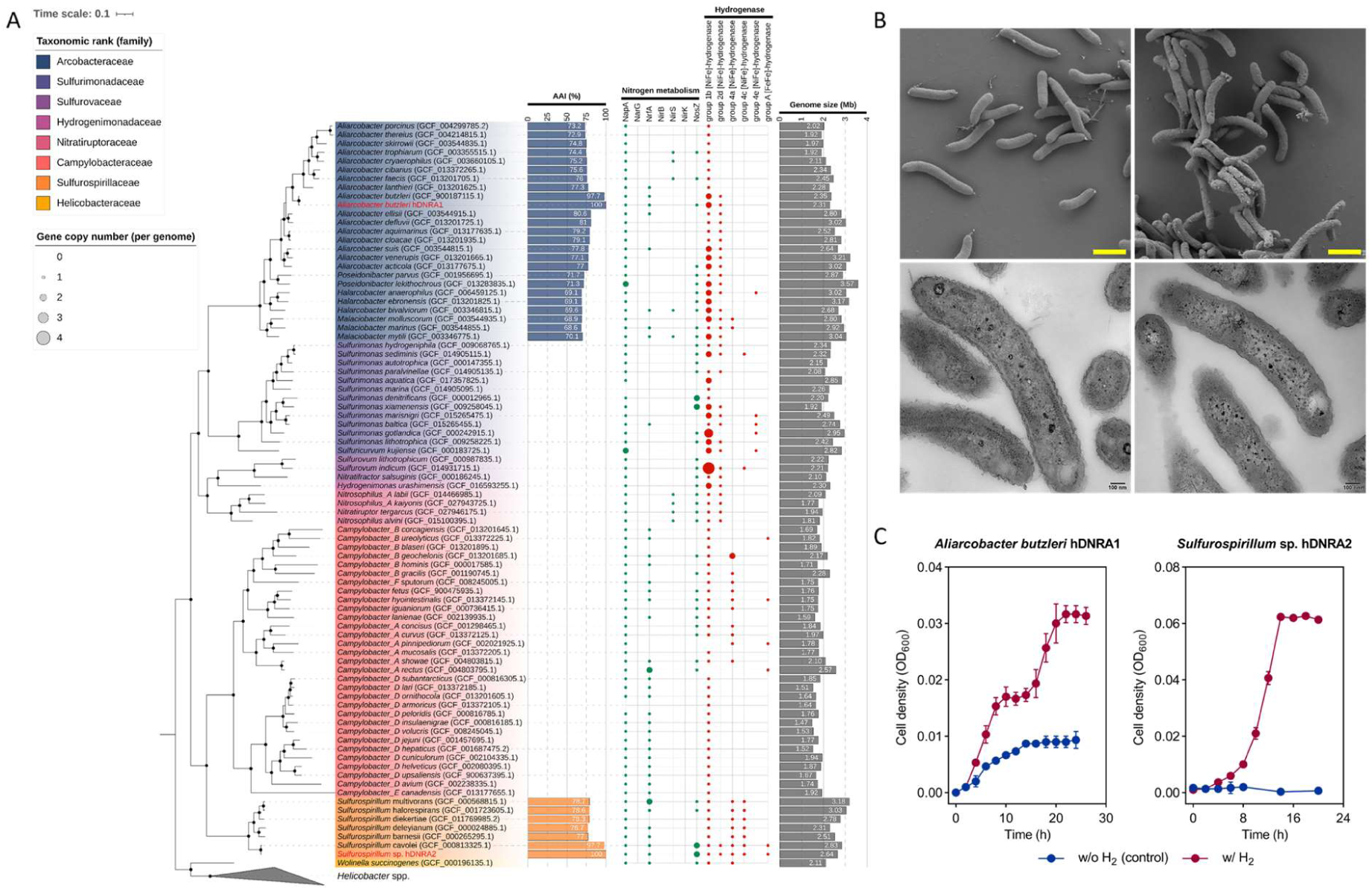
Phylogenetic affiliation and cellular morphology of *A*. *butzleri* hDNRA1 and *Sulfurospirillum* sp. hDNRA2 and their growth curves on hydrogenotrophic DNRA. (A) Maximum-likelihood phylogenetic tree of 92 *Campylobacterota* genomes generated using IQ-TREE v2.3.5, based on the alignment of the concatenated amino acid sequences of 120 single-copy bacterial housekeeping genes identified with GTDB-Tk v2.4.0 (release220). Seven *Helicobacter* spp. genomes served as the outgroup. Bootstrap values were derived from 10,000 bootstrap iterations. Bifurcations with >80% bootstrap values are marked with filled black circles. To the right of the tree is a grid plot visualizing the inventories of the genes encoding dissimilatory NO_3_^−^/NO_2_^−^ reductases and hydrogenases. (B) SEM (top) and TEM (bottom) images of *A*. *butzleri* hDNRA1 (left) and *Sulfurospirillum* sp. hDNRA2 (right). The scale bars, located at the bottom right corner of the micrographs, represent 1 µm and 0.1 µm on the SEM and TEM images, respectively. (C) Batch-culture growth curves for *A*. *butzleri* hDNRA1 and *Sulfurospirillum* sp. hDNRA2 provided with H_2_ as the electron donor and NO_3_^−^ as the electron acceptor. Each data point represents the average of three biological replicates (*n*=3), with the error bar representing the standard deviation.

A series of batch experiments performed with and without H_2_ (control) confirmed H_2_ as the sole electron donor for DNRA *sensu stricto* (that includes NH_4_ ^+^ production from NO_2_ ^−^) in these isolates. *A*. *butzleri* hDNRA1 exhibited growth on stoichiometric NO_3_ ^−^-to-NO_2_ ^−^ reduction in the absence of H_2_, with acetate as the probable electron donor and carbon source; however, NH_4_ ^+^ production from NO_2_ ^−^ reduction required H_2_ (Fig. 1C, 2A-C). In the presence of 5% (v/v) H_2_ in the headspace, *A*. *butzleri* hDNRA1 completely consumed 201 ± 1 µmol NO_3_ ^−^, of which 90 ± 2% (181 ± 2 µmol) was converted to NH_4_ ^+^, while exhibiting a diauxic growth. Oxidation of 670 ± 30 µmol H_2_ would have yielded 1340 ± 60 µmol e^−^, covering 84 ± 4% of the electron needed in dissimilatory nitrate ammonification (Eq. 1). Contrastingly, *Sulfurospirillum* sp. hDNRA2 was unable to couple NO_3_ ^−^ reduction with acetate oxidation (Fig. 2E). The electron yield from H_2_ oxidation (1,910 ± 20 µmol e^−^) greatly exceeded the theoretical electron demand for the observed NO_3−_- to-NH_4_ ^+^ reduction (1,330 ± 10 µmol e^−^), suggesting that *Sulfurospirillum* sp. hDNRA2 allocated ∼30% of the electrons from H_2_ oxidation for assimilation (Fig. 2F, G). With the same amount of NO_3−_ reduced to NH_4_ ^+^, *Sulfurospirillum* sp. hDNRA2 exhibited a significantly higher cell density (OD_600_ value of 0.064 ± 0.001 compared to 0.032 ± 0.002 of *A*. *butzleri* hDNRA1; *P* < 0.05) (Fig. 1C).

**Fig. 2.**
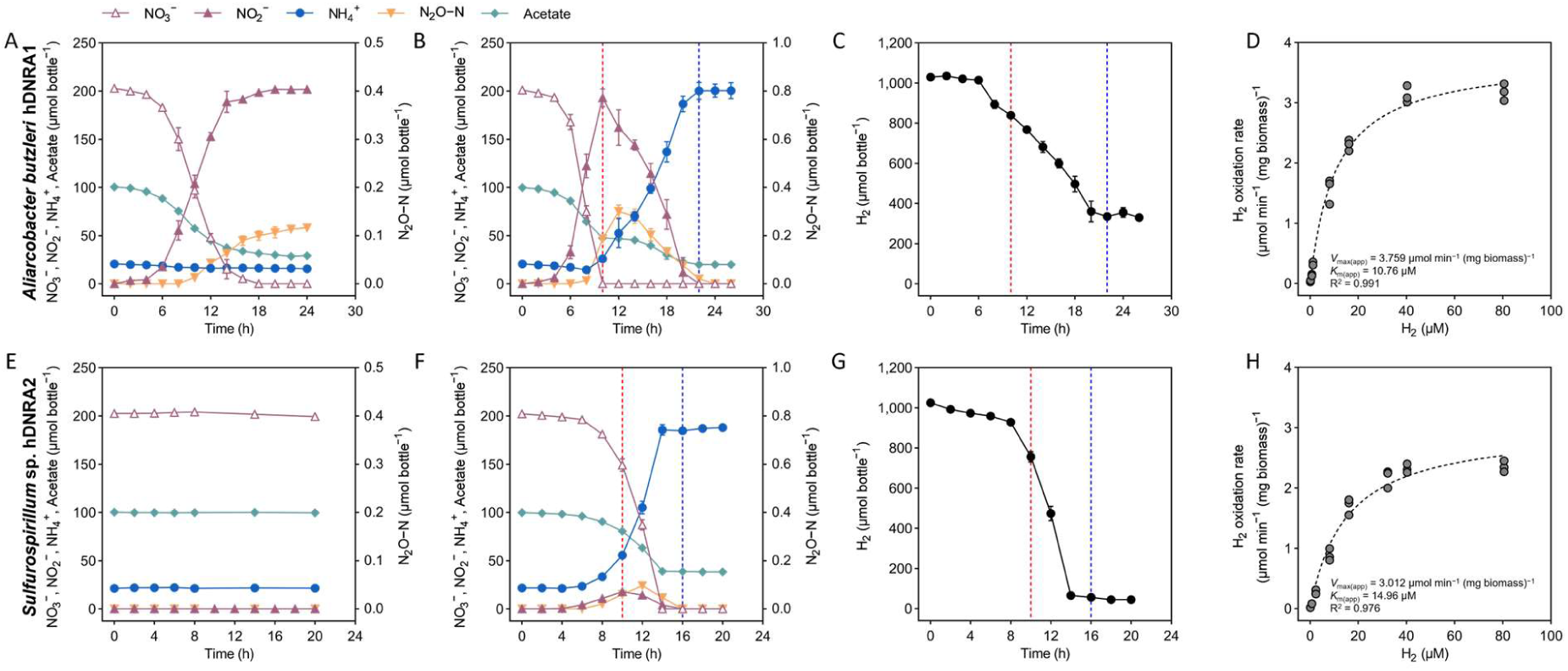
Hydrogenotrophic DNRA observed in batch cultures of *A*. *butzleri* hDNRA1 (A-C) and *Sulfurospirillum* sp. hDNRA2 (E-G). Reductive transformation of 2 mM NO ^−^ was monitored in batch cultures in the absence (A, E) and the presence of 5% (v/v) H_2_ (B, F) in the headspace. H_2_ consumption in the culture bottles was also monitored (C, G). The dotted vertical lines indicate the NO ^−^ peak (red) and NO_3−_/NO_2−_ depletion (blue). Each data point represents the mean of three biological replicates (*n*=3), with error bars indicating standard deviations. Additionally, the whole-cell Michaelis-Menten kinetics parameters of H_2_ oxidation are presented (D, H). Each point on the plot is the initial H_2_ oxidation rate determined from a batch incubation initiated with the molar concentration of dissolved H_2_ indicated on the *x*-axis.

Transient accumulation of the intermediates NO ^−^ and N O was observed in both isolates, but to a substantially larger degree in the *A*. *butzleri* hDNRA1 cultures. Neither isolate exhibited significant growth in the absence of acetate amendment; however, *Sulfurospirillum* sp. hDNRA2 exhibited significant NH_4_ ^+^ production (*P* < 0.05), which was sustained for at least 72 hours (Fig. S1). The *A*. *butzleri* hDNRA1 culture significantly reduced NO_3−_ to NO ^−^ (*P* < 0.05), but not to NH_4_ ^+^.

### H_2_ oxidation is directly coupled to DNRA

The dependence of DNRA *sensu stricto* on H_2_ in the two organisms was further verified through batch incubations with intermittent H_2_ starvation periods. As anticipated, *A*. *butzleri* hDNRA1 completely reduced 204 ± 4 µmol NO_3−_ before the first H_2_ starvation period, presumably with both acetate and H_2_ as electron donors (Fig. 3A). NO_2_ ^−^ turnover to NH_4+_ halted immediately after H_2_ was depleted. Each H_2_ injection (t = 34 and 59 h) was immediately followed by continuation of NO_2_ ^−^-to-NH_4_ ^+^ reduction. Acetate was consumed only when H_2_ was being consumed; however, a portion of electron from acetate may have been directed to NO_2_ ^−^ after repeated H_2_ starvation, as inferred from the discrepancy between the amounts of H_2_ oxidized (185 ± 6 µmol; 369 ± 13 µmol e^−^) and NO_2−_ reduced to NH_4_ ^+^ (96 ± 6 µmol; 578 ± 37 µmol e^−^) after the H_2_ addition at t = 59 h.

**Fig. 3.**
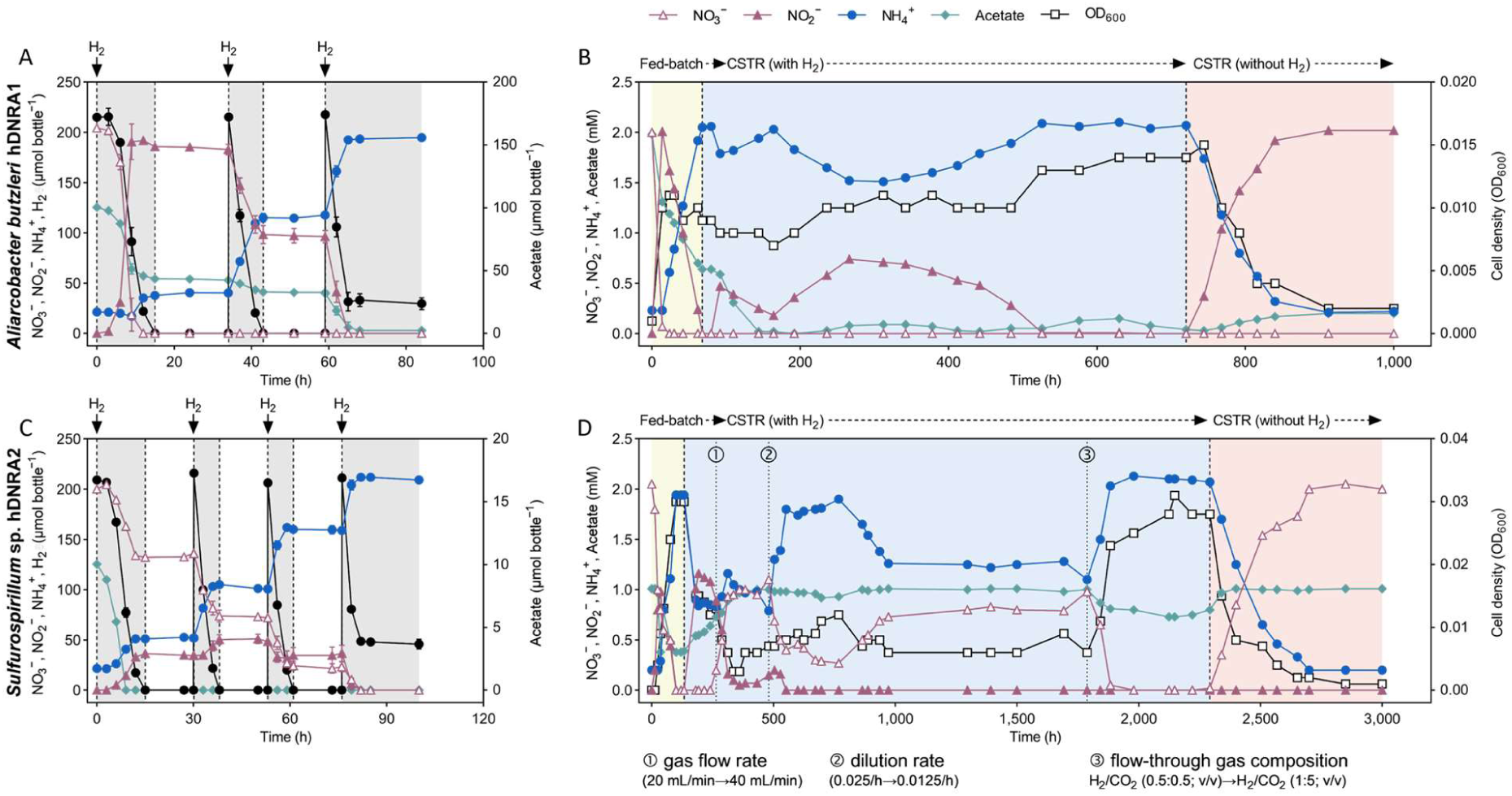
Coupling of H_2_ oxidation with DNRA examined in H_2_-limited batch cultures and chemostat reactor cultures of *A*. *butzleri* hDNRA1 (A, B) and *Sulfurospirillum* sp. hDNRA2 (C, D). The batch cultures were started with a stoichiometrically limiting amount of H_2_ (200 µmol bottle^−1^; a H_2_-to-NO_3_^−^ mol ratio of 1) and repeatedly replenished with 200 µmol bottle^−1^ H_2_ after confirmation of H_2_ depletion (indicated with black arrows). The concentrations of NO_3_^−^, NO_2_^−^, NH_4_^+^, H_2_ and acetate were monitored until the depletion of NO_3_^−^ and NO_2_^−^ and are presented as the total amount in the bottles. Each data point represents the mean of three biological replicates (*n*=3), with error bars indicating their standard deviations. Hydrogenotrophic DNRA by the two isolates was also examined in lab-scale chemostat reactors (B, D). In case of *Sulfurospirillum* sp. hDNRA2, the incubation conditions were optimized through the course of incubation to achieve a steady state, as indicated by the vertical dotted lines showing the time points at which the changes were applied with descriptions below. At the end of incubation H_2_ was removed from the gas stream.

The experiment with *Sulfurospirillum* sp. hDNRA2, performed identically but with a smaller initial amount of acetate (10 µmol bottle^−1^), not only confirmed that H_2_ served as the sole electron donor for both NO_3−_- to-NO_2_ ^−^ and NO_2_ ^−^-to-NH_4_ ^+^ reduction, but also showed that these reduction reactions could be sustained without organic carbon source for an extended period (Fig. 3C). Also notably, NO_3_ ^−^-to-NO_2_ ^−^ and NO_2_ ^−^-to- NH ^+^ reduction occurred simultaneously. The ratio of electron distribution to NO_3_ ^−^ and NO_2_ ^−^ from the initially added H_2_ was 0.70. This ratio dropped to 0.44 for H_2_ added at t = 30 h and further decreased to 0.28 for H_2_ added at t = 53 h, indicating a shift in electron allocation between NO_3−_ and NO_2−_. Even with halted cell growth after acetate depletion at t = 9 h, DNRA activity was sustained for >70 hours, completely reducing 163 ± 1 µmoles NO_3−_ and 14.5 ± 1.1 µmol NO_2_ ^−^ to 173 ± 3 µmol NH_4_ ^+^.

The whole-cell kinetics of H_2_ oxidation coupled to NO_2_ ^−^-to-NH_4_ ^+^ reduction by both *A*. *butzleri* hDNRA1 and *Sulfurospirillum* sp. hDNRA2 fitted well to the Michaelis-Menten kinetics model (R^2^ > 0.97; Fig. 2D, H). The *K*_m(app)_ values were calculated to be 10.8 µM (95% confidence interval: 9.0-12.9) and 15.0 µM (11.5-19.5) for *A*. *butzleri* hDNRA1 and *Sulfurospirillum* sp. hDNRA2, respectively. The maximum H_2_ oxidation rates (*V*_max(app)_) were 3.76 µmol min^−1^ (mg biomass)^−1^ (3.56-3.97) for *A*. *butzleri* hDNRA1 and 3.01 µmol min^−1^ (mg biomass)^−1^ (2.76-3.30) for *Sulfurospirillum* sp. hDNRA2. Therefore, the two isolates share a comparable target H_2_ concentration range and maximum turnover rates.

### Hydrogenotrophic DNRA is sustained during continuous cultivation

The feasibility of establishing a steady state culture with the two isolates was examined, as this is often considered as a prerequisite for harnessing microbial metabolisms for environmental applications (Fig. 3B, D). The chemostat culture of *A*. *butzleri* hDNRA1 successfully reached a steady state after 526 hours, with NO_3−_ in the influent being completely reduced to NH_4_ ^+^ (2.07 ± 0.02 mM) in the reactor supplied with a continuous stream of H_2_/CO_2_/N_2_ (0.5:0.5:99) mixed gas (Fig. 3B). At steady state, the rates of NO_3_ ^−^ consumption and NH_4_ ^+^ production were 30.5 ± 0.6 µmol h^−1^ and 28.1 ± 0.4 µmol h^−1^, respectively. Acetate was consumed at 13.7 ± 0.7 µmol h^−1^, with its steady state concentration sustained above zero (0.09 ± 0.05 mM, versus 1 mM in the influent). H_2_ concentrations in the influent and effluent gas streams were indistinguishable (*P* > 0.05), consistent with stoichiometric calculations showing H_2_ was supplied in excess (4,018 μmol h^−1^ in the flow- through gas, compared to the theoretical demand of 245 μmol h^−1^, assuming H_2_ serves as the sole electron donor for NO_3−_-to-NH_4_ ^+^ reduction). After observing the steady-state reactor operation for approximately 200 hours (i.e., the time length equivalent to flow-through of five reactor volumes), the gas supply was switched to exclude H_2_ and the NO_2−_ concentration in the reactor gradually increased. A new steady state was established, at which the NO_2_ ^−^ concentration in the reactor matched stoichiometrically with the NO_3_ ^−^ in the influent, corroborating that the DNRA *sensu stricto* was dependent on the oxidation of H_2_, but not acetate.

Establishing a *Sulfurospirillum* sp. hDNRA2 continuous culture required rigorous parameter optimization (Fig. 3D). The initial condition failed to achieve the targeted steady state with 2 mM NO_3_ ^−^ being completely transformed to NH_4+_. Only after H_2_ and CO_2_ concentrations in the flow-through gas were increased to 1% and 5%, respectively, the reactor completely reduced 2 mM NO_3_ ^−^ to NH_4_ ^+^ at a rate of 15.65 ± 0.24 µmol h^−1^. During this steady-state operation, acetate was consumed at 1.73 ± 0.35 µmol h^−1^. Exclusion of H_2_ from the influent gas led to immediate increase in NO_3−_ concentration and decrease in NH_4_ ^+^ concentration eventually resulting in reactor washout, which clearly demonstrated the physiological difference between the two isolates, in that reduction of NO_3_ ^−^, as well as NO_2_ ^−^, was dependent on H_2_ in *Sulfurospirillum* sp. hDNRA2.

### Both strains encode hydrogenases and Nap and Nrf for hydrogenotrophic DNRA

Closed genomes of *A. butzleri* hDNRA1 and *Sulfurospirillum* sp. hDNRA2 were obtained through hybrid assembly of short-read and long-read data (Sequencing statistics in Table S1). Both strains possess a *nap* cluster and a *nrf* cluster encoding the respiratory periplasmic nitrate reductase (Nap; *napAGHBFLD*) and an ammonia-forming cytochrome *c*_552_ nitrite reductase (Nrf; *nrfAH*), respectively (Fig. 4A and Table S2). The *nrf* cluster is immediately downstream of the *nap* cluster in *A*. *butzleri* hDNRA1, and in close proximity (∼30 kb apart) in *Sulfurospirillum* sp. hDNRA2, corroborating their functional relatedness. Interestingly, although neither genome contained *nirK* or *nirS*, confirming the inability to denitrify, both genomes harbor clade II *nosZ* genes encoding the catalytic subunit of nitrous oxide reductase (Nos).

**Fig. 4.**
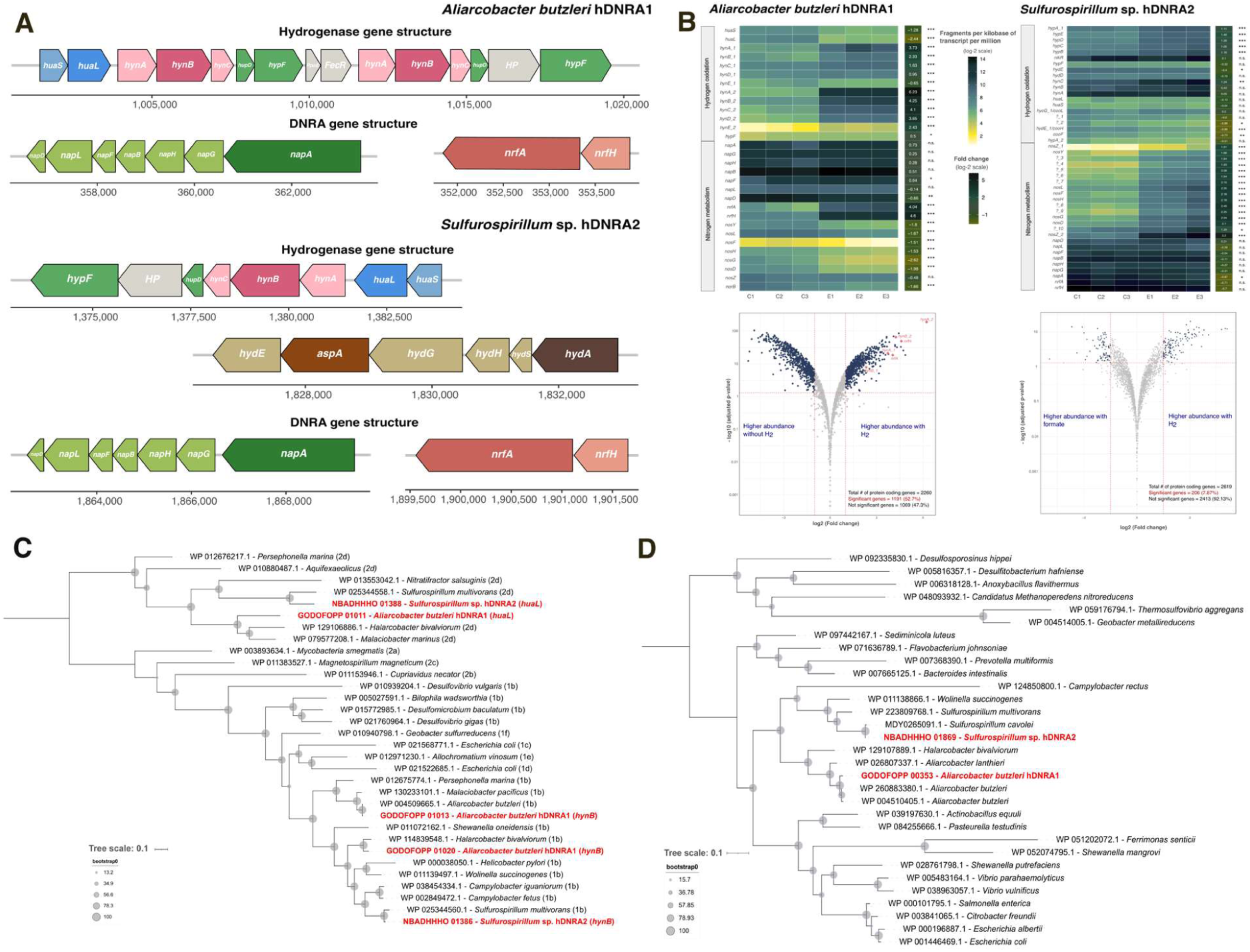
Genomic and transcriptomic bases for hydrogenotrophic DNRA in *A*. *butzleri* hDNRA1 and *Sulfurospirillum* sp. hDNRA2. (A) Clusters of the genes putatively associated with hydrogenotrophic DNRA identified from the complete genomes of *A*. *butzleri* hDNRA1 and *Sulfurospirillum* sp. hDNRA2. (B) Differential expression analyses of the functional genes involved with hydrogen oxidation and nitrogen metabolism (top), and genome-wide transcription changes presented in volcano plots (bottom), comparing the cultures grown under H_2_-free (acetate and formate provided as the alternate electron donor for *A*. *butzleri* hDNRA1 and *Sulfurospirillum* sp. hDNRA2, respectively) versus H_2_-supplemented conditions. Additionally, maximum-likelihood phylogenetic trees generated with the amino acid sequences of the catalytic subunits of the group 1b [NiFe]- and group 2d [NiFe]-hydrogenases (C) and the cytochrome *c*_552_ nitrite reductase (D) are presented to illustrate the phylogenetic positioning of these key functional genes found in the two hydrogenotrophic DNRA isolates.

Structural and maturation genes for multiple hydrogenases were also identified in the genomes. Both bacteria co-encode a group 1b [NiFe]-hydrogenase (*hynAB*) with a group 2d [NiFe]-hydrogenase (*huaSL*).

Additionally, in the *A*. *butzleri* hDNRA1 genome, another set of *hynAB* genes with 51.2% amino acid identity, was identified in the vicinity. Both isolates also possessed the *hypABCDEF* genes required for the maturation of [NiFe]-hydrogenases. Additionally, *Sulfurospirillum* sp. hDNRA2 harbored group 4a and 4c [NiFe]-hydrogenases to couple the oxidation of formate and carbon monoxide to H_2_ generation from protons, as well as a group A [FeFe]-hydrogenase flanked by H-cluster maturation assembly genes (*hydGEF*) and aspartate ammonia-lyase (*aspA*) (Table S3). This suggests this strain can mediate a range of processes dependent on H_2_ oxidation and production, in addition to hydrogenotrophic DNRA. In addition to its formate hydrogenlyase complex, this strain also encodes a formate dehydrogenase H (*fdhF*), which may explain its ability to utilize formate as an electron donor (Fig. S2).

The capacity for mixotrophic growth, specifically biomass production through carbon acquisition from acetate and CO_2_, was also evident from the *Sulfurospirillum* sp. hDNRA2 genome (Fig. S3). Acetate assimilation likely depends on the acetate kinase and phosphate acetyltransferase encoded by *ackA* and *pta* genes, respectively, which are present also in *A*. *butzleri* hDNRA1. In *Sulfurospirillum* sp. hDNRA2, acetyl- CoA may also be generated by the activity of acetate-CoA ligase encoded by the *acs* gene. The reductive tricarboxylic acid (rTCA) cycle is the most likely pathway employed by *Sulfurospirillum* sp. hDNRA2 to assimilate CO_2_, as many of the genes encoding this pathway, including those encoding 2-oxoglutarate synthase, isocitrate dehydrogenase, pyruvate carboxylase, and succinate-CoA ligase (ADP-forming), as well as pyruvate synthase that catalyzes the carboxylation of acetyl-CoA, are present in the genome. However, two of the genes encoding key enzymes in the conventional rTCA cycle, phosphoenolpyruvate carboxylase and ATP citrate lyase, are missing from the genome. Involvement of the Wood-Ljungdahl pathway for CO_2_ fixation is unlikely, as the gene encoding a pivotal enzyme for the pathway, CO- methylating acetyl-CoA synthase, was not identified in the genome. The lack of many of these putative rTCA cycle genes and the absence of CO-methylating acetyl-CoA synthase in *A. butzleri* hDNRA1 explains its dependence on acetate for biomass carbon.

### Transcriptomic underpinning of hydrogenotrophic DNRA

Differentially expressed gene (DEG) analysis revealed significant differences between the transcriptomes of *A*. *butzleri* hDNRA1 cultures grown with H_2_ (coupled with NO_2_ ^−^-to-NH_4_ ^+^ reduction) and those grown without H_2_ (NO_3_ ^−^-to-NO_2_ ^−^ reduction coupled to acetate oxidation) (Fig. 4B). Statistically significant and meaningful differences (|log_2_ (fold change)| > 1 and *P* < 0.05) were observed for 1,191 out of 2,263 protein-coding genes, with 547 genes up- regulated under the H_2_-oxidizing condition. The high transcript levels and significant upregulation of the two sets of *hynAB* genes (3.7 ± 0.4-/2.3 ± 0.4- and 6.2 ± 0.2-/4.2 ± 0.2 fold; *P* < 0.001) and the *nrfA* gene (4.0 ± 0.4-fold; *P* < 0.001) in the presence of H_2_ suggested that the group 1b [NiFe]-hydrogenases and the cytochrome *c*_552_ nitrite reductase catalyze H_2_ oxidation and NO_2−_-to-NH_4_ ^+^ reduction, respectively. The transcription of the *napAGHBFLD* cluster was not significantly affected by the presence of H_2_, except for *napF* (up-regulated; *P* < 0.05) and *napD* (down-regulated; *P* < 0.01). The transcription of *napA* was at least 6.5-fold higher than that of the single-copy housekeeping genes *dnaA*, *rho*, and *recA*, respectively, regardless of the presence of H_2_, corroborating that NO_3_ ^−^-to-NO_2−_ reduction is mediated by the NapA. The downregulation of the *huaS* and *huaL* genes in the presence of H_2_ suggests that these genes are unlikely to participate in hydrogenotrophic DNRA.

For *Sulfurospirillum* sp. hDNRA2, transcriptome analysis focused on identifying genes that were highly expressed during hydrogenotrophic DNRA relative to single-copy housekeeping genes, given no DNRA- or H_2_-independent growth conditions have yet been identified for this organism. Among the 2,619 protein-coding genes, > 344 genes (13.2%), including those encoding catalytic subunits of respiratory hydrogenases and DNRA-catalyzing enzymes, exhibited higher transcription levels than all scrutinized single-copy housekeeping genes (Fig. S4). Both *hynAB* genes showed transcription levels at least four times higher than that of *recA*. The functional genes encoding the DNRA pathway, i.e., *napA* and *nrfA*, were among the top 1% most highly transcribed genes. Interestingly, one of the duplicate *nosZ* copies in the genome was also one of the 10 most highly differentially expressed genes. Additionally, genes encoding enzymes of the rTCA cycle, such as *sucCD* (succinate-CoA ligase), *gltA* (citrate synthase), *korAB* (2-oxoglutarate synthase) and *porA* (pyruvate synthase), exhibited higher transcription levels than *recA* (Fig. S5).

## DISCUSSION

Thermodynamically and biochemically, the redox coupling of H_2_ oxidation and DNRA is not unexpected, as the combined reaction is highly exergonic, and various configurations of electron transport chains can bridge the substantial redox potential difference (Δ*E*_0_′ = 0.76 V) between the two half-reactions (10, 30). Insinuations of hydrogenotrophic DNRA, in fact, abound in the literature. Several publications reporting the discovery and characterization of new bacterial isolates, including *Desulfurobacterium indicum* and *Ammonifex degensii*, denoted the ability of these isolates to utilize H_2_ as an electron donor and NO_3_ ^−^/NO_2_ ^−^ as an electron acceptor (31, 32). Additionally, several genomes (including metagenome-assembled genomes) possessing hydrogenase genes (e.g., *hynAB*) and *nrfA* have been reported (33, 34). In these studies, however, the capability of these isolates to couple H_2_ oxidation with DNRA for energy conservation was not examined. A recent study on the physiology of *Trichlobacter ammonificans*, a newly isolated *Geobacter*-like organism with an octaheme cytochrome *c* nitrite reductase (a distant relative of pentaheme NrfA), hinted that H_2_ may serve as a supplemental electron donor for NO_2_ ^−^-to-NH_4_ ^+^ reduction; however, this organism required acetate as the primary electron donor, and the contribution of H_2_ as an electron source could not be separated from that of acetate (35). The significance of the current discovery lies in that the physiological dependency of DNRA *sensu stricto* on H_2_ presence was unequivocally verified in two distinct, genomically characterized isolates. The reaction stoichiometry indicates a tight redox coupling between H_2_ oxidation and DNRA, further supported by genome and transcriptome data implicating group 1b [NiFe]-hydrogenases (*hynAB*) and cytochrome *c*_552_ nitrite reductases (*nrfAH*) in this novel pathway (36, 37).

The dependency of DNRA on electrons donated from H_2_, rather than organic carbon, was evident. For both organisms, NH_4_ ^+^ production from NO_2_ ^−^ reduction required H_2_ presence and was accompanied by a near-stoichiometric H_2_ consumption. Naturally, as organic carbon (acetate) was necessary only for biomass synthesis, hydrogenotrophic DNRA was independent of the C:N ratio and redox potential, contrary to the generally perceived hypothesis with regards to the regulation of the DNRA pathway (15, 19, 38). Hydrogenotrophic DNRA in *Sulfurillospirillum* sp. hDNRA2 occurred even in the absence of dissolved organic carbon (Fig. 3C; see t > 70 h), where the low ratios of electron donor (H_2_) to electron acceptor (NO_3_ ^−^ or NO_2_ ^−^) defined highly oxidizing conditions, typically considered unsuitable for DNRA. This observation suggests that the competition between DNRA and denitrification determining NO_3_ ^−^ fate, traditionally explained by the C:N ratio and redox potential, may demand a revision for ecological contexts where H_2_ is available via *in situ* production or transport (39). The competition between hydrogenotrophic DNRA and denitrification in such ecological settings may also be an interesting subject of future research, for a more comprehensive understanding of environmental nitrogen cycling (40).

The two isolates, despite being within the same phylum (89.1% 16S rRNA gene identity and 69.0% ANI) and putatively sharing the same set of enzymes for H_2_ oxidation (encoded by *hynAB*) and DNRA (encoded by *napA* and *nrfA*), exhibited several notable physiological differences. They differed in their ability to utilize acetate as the electron donor for NO_3_ ^−^-to-NO_2_ ^−^ reduction, and also in their capacity to carry out hydrogenotrophic DNRA without acetate as the carbon source. The extent of NO_2_ ^−^ accumulation during NO_3_ ^−^-to-NH_4_ ^+^ reduction was also different, suggesting that hydrogenotrophic DNRA in *Sulfurillospirillum* sp. hDNRA2 was not inhibited by the NO_3−_, unlike *A*. *butzleri* hDNRA1 or the previously isolated organotrophic DNRA-catalyzing bacteria (16, 41). These subtle but important physiological differences cannot be predicted through analyses of their genomes, emphasizing the isolation of new microorganisms and experimental interrogation of their physiology still remains to be vital for furthering our understanding of the biogeochemical processes and grasping the endless biotechnological potentials of microorganisms and microbiota in the nature (42). Securing more diverse hydrogenotrophic DNRA isolates may lead to the discovery of surprising physiological diversity with wide-reaching environmental implications and utility, such as fully chemolithoautotrophic DNRA or DNRA coupling with high-affinity, and even atmospheric, H_2_ oxidation (43, 44).

Hydrogenotrophic DNRA metabolism holds promising biotechnological potential. Today, the environmental consequences of nitrogen fertilization are of great concern, as many freshwater and coastal ecosystems across the globe suffer from severe eutrophication, and greenhouse gas emissions associated with the production and application of nitrogen fertilizers account for a substantial portion of global emissions (45, 46). A substantial portion of nitrogenous fertilizers, mostly applied to agricultural soils in the forms of NH_4_ ^+^ and organic nitrogen (e.g., urea), is microbially oxidized to the more mobile form, NO_3_ ^−^, and subsequently lost to freshwater bodies via runoff and leaching or denitrified to gaseous forms, causing nitrogen loss and diminishing nitrogen use efficiency (47). Diminished fertilizer use efficiency is especially problematic, as nitrogen fertilizer production, still relying entirely on the Harber-Bosch process, is an energy- and carbon-intensive process, and the microbial transformation processes involved, i.e., nitrification and denitrification, emit N_2_O (5, 48). Hydrogenotrophic DNRA offers a novel biotechnological opportunity to address this nitrogen dilemma by utilizing minimal amounts of green hydrogen generated *in situ* to convert the freshwater contaminant, NO_3_ ^−^, into a fertilizer substitute, NH_4_ ^+^ (the schematic shown in Fig. S6). The current study demonstrated that stable continuous operation of H_2_-fed DNRA chemostat was feasible, with provision of 4 mmol h^−1^ H_2_ at a modest 0.5% concentration sufficient to completely convert 48 μmol h^−1^ NO_3_ ^−^ to NH_4_ ^+^. Sourcing such a small flux of H_2_ would not be challenging; however, much remains to be examined for properly assessing the practical feasibility of this conceptual biotechnology. Consistent influx of dissolved oxygen and competitive challenges from inflowing denitrifiers will be unavoidable in the reactor, and engineering solutions to sustain hydrogenotrophic DNRA activity under such adversities may need to be devised. If technically and economically viable, this promising biotechnology can simultaneously address two of the most pressing sustainability issues: climate change and the disruption of biogeochemical cycles (4).

## MATERIALS AND METHODS

### Culture medium and growth conditions

The defined medium used for enrichment, isolation, and routine cultivation of hydrogenotrophic DNRA bacteria contained, per liter: 4.76 mmol NaCl, 0.47 mmol CaCl_2_·2H_2_O, 0.24 mmol MgCl_2_·6H_2_O, 0.2 mmol NH_4_Cl, and 1 mL of a 1000× modified Wolin’s mineral solution (DSMZ medium 141). Batch incubations were conducted with an initial aqueous-phase volume of 100 mL in 590-mL glass bottles, which were sealed with bromobutyl-rubber stoppers fitted into GL45 open-topped screw caps. After autoclaving (121 °C for 20 min), a filter-sterilized PIPES (500 mM; pH 7.0) was added to a final concentration of 5 mM. The medium was further supplemented with filter-sterilized 200× Wolin’s vitamin solution (DSMZ medium 141). Bottles were equilibrated with a pressurized 5% H_2_/5% CO_2_/90% N_2_ (World-Enersys, Daejeon, South Korea), unless otherwise specified. Equilibration with the CO_2_-containing gas mixture reduced the final pH to 6.5 ± 0.1. Immediately before inoculation, sodium acetate and KNO_3_ were added to target concentrations from sterile stock solutions. Agar plates were prepared by adding 1.5% (w/v) Bacto agar (BD, Franklin Lakes, NJ) to the liquid medium. Inoculated agar plates were incubated in an anaerobic chamber (Coy Laboratory Products, Inc., Grass Lake, MI) with an internal atmosphere of 4% H_2_/5% CO_2_/91% N_2_.

### Isolation, screening, and cultivation

Activated sludge for isolating hydrogenotrophic DNRA-catalyzing bacteria was collected from the anoxic segment of the municipal wastewater treatment plant (WWTP) in Daejeon, South Korea (36°23’09.4”N 127°24’28.5”E) on October 26, 2020. The sample was promptly transported in a 1-L polyethylene bag kept in a cooler and stored at 4 °C in the dark upon arrival at the laboratory. Initially, the goal was to isolate DNRA-catalyzing microorganisms that utilize acetate as the sole electron donor. In an anaerobic chamber, 20 mL activated sludge was diluted into 200 mL autoclaved medium with 10 mM acetate and 2 mM NO_3_ ^−^ in a loosely capped 590-mL glass bottle. To inhibit fungal growth, cycloheximide (Sigma-Aldrich, St. Louis, MO) was added to a concentration of 0.01% (w/v). The enrichment culture was stirred with a magnetic bar at 400-500 rpm in the dark. Every two weeks, 20 mL of the enrichment culture was transferred to 200 mL of fresh medium. After three transfers, the enrichment culture was serially diluted in autoclaved medium and spread onto agar plates containing 20 mM acetate and 10 mM NO_3_ ^−^. Bacterial isolates from single colonies were then subjected to DNRA phenotype screening (16).

### Discovery, verification, and physiological characterization of hydrogenotrophic DNRA in the *Campylobacterota* isolates

The DNRA activity of two screened isolates, later identified as belonging to the genera *Aliarcobacter* and *Sulfurospirillum*, was investigated using batch cultures. Initially, the 200-µL cell suspensions prepared from single colonies were transferred into 100-mL medium containing 2 mM NO_3_ ^−^ and 10 mM acetate, then equilibrated with N_2_. After failing to reproduce DNRA activity in sealed batch cultures, the possibility of H_2_, a component of the anaerobic chamber atmosphere, serving as the electron donor was examined. Three sets of batch cultures were prepared: (1) 5% H_2_/5% CO_2_/ 90% N_2_ headspace and 10 mM acetate in the medium, (2) 5% H_2_/95% N_2_ headspace and 10 mM acetate in the medium, and (3) 5% H_2_/5% CO_2_/90% N_2_ headspace with no organic carbon. Additionally, a set of H_2_-free controls was prepared with 95% N_2_/5% CO_2_ headspace and 10 mM acetate in the medium. All cultures were prepared with 2 mM NO_3_ ^−^. The H and N O concentrations in the headspace, the dissolved concentrations of NO_3−_, NO ^−^, NH_4_ ^+^, and acetate, and the cell density (OD_600_) were monitored until no further change was observed. The pH was measured at the end of each incubation to ensure that no significant shift occurred.

The H_2_-dependency of DNRA in *A*. *butzleri* hDNRA1 and *Sulfurospirillum* sp. hDNRA2 was further verified in the experiment with repeated addition of limiting amounts of H_2_. The initial concentration of NO_3-_ was 2 mM, and the headspace composition consisted of 5% CO_2_ and 95% N_2_ before H_2_ addition. The initial acetate concentration was 1 mM for the *A*. *butzleri* hDNRA1 cultures and 0.1 mM for the *Sulfurospirillum* sp. hDNRA2 cultures. A limiting amount (200 μmoles) of H_2_ was initially injected to the headspace. The dissolved concentrations of NO_3−_, NO ^−^, and NH_4_ ^+^, and the headspace H_2_ concentration were monitored throughout incubation, and upon each H_2_ depletion, 200 μmoles H_2_ was injected, following confirmation of discontinued NO_3_ ^−^/NO_2_ ^−^ reduction.

### Determination of whole-cell Michaelis-Menten kinetics of anaerobic H_2_ oxidation

The whole-cell Michaelis-Menten kinetics parameters, *V*_max(app)_ and *K*_m(app)_, were experimentally determined to assess the ability of *A*. *butzleri* hDNRA1 and *Sulfurospirillum* sp. hDNRA2 to utilize varying concentrations of H_2_ as an electron donor for NO_2_ ^−^-to-NH_4_ ^+^ reduction. Each experiment was performed with cultures grown with 1 mM acetate and 2 mM NO_3_ ^−^ in the medium and with 5 % H_2_ and 5% CO_2_ in the headspace. Immediately after confirming depletion of NO_3_ ^−^ and NO_2_ ^−^, culture bottles underwent three vacuuming-and-pressurizing cycles with N₂ gas to remove any residual H₂, followed by relief of excess pressure. The dry biomass was measured with triplicate 100-mL cultures grown under identical conditions and harvested at the same cell density using the previously described method (49). NO₂⁻ was added to a concentration of 10 mM, and H₂, generated using a PG-H₂ Plus Hydrogen Generator (PerkinElmer, Inc., Waltham, MA), was added to the culture bottles using a 1700 series Gastight syringe (Hamilton Company, Reno, NV) to target concentrations ranging between 100 and 100,000 ppmv. The initial linear H_2_ consumption rate was calculated from headspace H_2_ concentrations measured at 20-minute intervals. A nonlinear least-squares regression analysis was performed using GraphPad Prism v9.5.1 to estimate *V*_max(app)_ and *K*_m(app)_ from the dataset of H_2_ consumption rates versus dissolved H_2_ concentrations, collected from triplicate experiments.

### Continuous anaerobic incubation of *Campylobacterota* isolates on hydrogenotrophic DNRA

A chemostat reactor was set up with a custom-designed glass vessel with a total capacity of 1.14-L and a working volume of 600 mL (Fig. S7). The main reactor vessel was fed at a dilution rate of 0.025 h^−1^ with fresh medium containing 2 mM NO ^−^ and 1 mM acetate from a 5-L glass bottle, maintained anoxic with a continuous N_2_ stream (20 mL min^−1^) and stirred with a magnetic bar at 600-700 rpm. The culture was started as a batch culture continuously supplied with a 30 mL min^−1^ stream of filter-sterilized mixed gas composed of 0.5% H_2_/0.5% CO_2_/99% N_2_. The reactor was transitioned to a chemostat immediately after NO_3_ ^−^/NO_2_ ^−^ were depleted. The dissolved NO_3_ ^−^, NO_2_ ^−^, and NH_4_ ^+^ concentrations and cell density were monitored throughout the reactor operation. As for *Sulfurospirillum* sp. hDNRA2, several adjustments to the mixing ratio and the flow rate of the gas stream were made during the incubation. Once the concentrations of the N species were stabilized for >200 hours, indicating a steady state, H_2_ was excluded from the mixed gas stream to confirm the H_2_ dependency of NH_4_ ^+^ production from NO_3−_.

### Analytical methods

H_2_ concentrations in the headspace were measured using an 8890 gas chromatography (GC) system equipped with a thermal conductivity detector and serially connected packed MolSieve 5Å (1.83 m × 2.0 mm, 60/80 mesh) and Porapak Q (1.83 m × 2.0 mm, 80/100 mesh) columns (Agilent Technologies, Inc., Santa Clara, CA). Ar (≥99.999%) was used as the carrier and reference gas. The inlet and detector temperatures were set to 150 °C and 230 °C, respectively, and the oven temperature was ramped up to 150 °C at a constant rate of 25 °C min^−1^ after being held constant at 40 °C for 3 min. N_2_O concentrations were determined using a 8890 gas chromatograph equipped with a micro-electron capture detector and a HP-PLOT Q capillary column (30 m × 0.32 mm, 20 µm film thickness). He (≥99.999%) was used as the carrier gas and 5% CH_4_/ 95% Ar gas as the make-up gas. The inlet, oven, and detector temperatures were set to 200 °C, 85 °C, and 250 °C, respectively. Gas samples were manually injected using 1700 series Gastight syringes. The dissolved concentrations of H_2_ and N_2_O were calculated from the measured headspace concentrations using dimensionless Henry’s constants (aqueous/gaseous; 25 °C) of 0.0193 for H_2_ and 0.595 for N_2_O (50). The concentrations of NO_3_ ^−^, NO_2−_, and NH_4+_ were determined colorimetrically (51, 52). Acetate concentration was determined using a Prominence high-performance liquid chromatography (HPLC) system equipped with a UV-VIS detector (Shimadzu, Kyoto, Japan) and an Aminex HPX-87H column (300 mm × 7.8 mm; Bio-Rad Laboratories Inc., Hercules, CA).

### DNA extraction and genome sequencing

Genomic DNA was extracted from batch cultures of *A*. *butzleri* hDNRA1 and *Sulfurospirillum* sp. hDNRA2 using DNeasy blood & tissue kit (Qiagen, Germany). A hybrid genome sequencing approach was employed to obtain closed genomes, combining HiSeq (Illumina, Inc., San Diego, CA) short-read sequencing with MinION (Oxford Nanopore Technologies, Oxford, UK) long- read sequencing. For short-read sequencing, a paired-end sequencing library was prepared using TruSeq DNA PCR-Free kit (Illumina, Inc.), and the library was sequenced on a Hiseq X Ten platform at Macrogen, Inc. (Seoul, South Korea) with a targeted throughput of 4 Gb. Raw reads were quality-trimmed using Trimmomatic v0.36, and *de novo* assembly was performed using SPAdes v3.15.3 with a contig size cutoff of 200 bp (53, 54). Long-read sequencing was performed using a MinION sequencer with a R9.4.1 flowcell. The sequencing library was prepared with Ligation Sequencing kit (Oxford Nanopore Technologies, Oxford, UK), and base-calling and demultiplexing were performed with Guppy v6.2.11. Adapter sequences were trimmed using Porechop v0.2.4 (55). The two genome sequence datasets were merged using Unicycler v0.4.8 (‵conservative′ mode) to construct a short-read-first hybrid assembly (56). The completed genomes were annotated using Prokka v1.14.5 (57).

### RNA extraction and transcriptome sequencing

For RNA extraction, triplicate cultures were incubated in a 5-L glass bottle containing 1 L of medium supplemented with 1 mM acetate and 2 mM NO_3_ ^−^ and 5% (v/v) H_2_ in the headspace. Additionally, the cultures of *A*. *butzleri* hDNRA1 grown without H_2_ and *Sulfurillosprillum* sp. hDNRA2 grown without H_2_, but with 10 mM formate were also prepared. Cell density, H_2_, and NO_3_ ^−^/NO_2_ ^−^ concentrations were monitored to ensure that the samples for transcriptome were collected during the exponential phase (Fig. S8). The entire batch was filtered through a 0.22-µm Sterivex filter unit (Merck Millipore, Germany). Subsequently, 10 mL RNAprotect bacteria reagent (Qiagen) was passed through the filter, and then stored at −80 °C. After thawing on ice, the membrane filter was aseptically separated from the filter unit and disrupted using acid-washed glass beads. The resulting crude lysate was processed sequentially with the RNeasy Mini kit, RNase-Free DNase and the RNeasy MinElute cleanup kit (Qiagen). Following treatment with the Ribo-Zero Plus rRNA depletion kit, the RNA-Seq library was prepared using TruSeq Stranded Total RNA library prep kit (Illumina). Paired-end (101 bp × 2) sequencing was performed on a NovaSeq 6000 platform at Macrogen, Inc.

### Transcriptome analyses

The raw reads were quality-trimmed using Trimmomatic v0.36 with default parameters and mapped to the genomes using Bowtie2 v2.5.1 (58). The alignments in SAM format were converted to BAM format and sorted using SAMtools v1.18 (59). Read counts for each gene were obtained with featureCounts v2.0.6 (60). The FPKM (fragments per kilobase of transcript per million reads mapped) values for normalized quantification of the targeted genes were computed using equation 3.

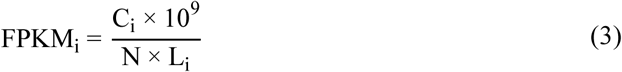

where *C_i_* is the count of fragments mapped onto the gene of interest, *N* is the total number of fragments mapped per kilobase of the genome, and *L_i_* is the length of the gene of interest in kilobases. The analyses of the transcription data were performed using the ’*DESeq2’* package v1.42.0, with the transcriptome data from biological triplicates as inputs (61). In each pairwise comparison, a criterion of |log_2_ (fold change)| > 1 and *P <* 0.05 was used to identify differentially expressed genes.

### Statistical analysis

All experiments except the chemostats were conducted in biological triplicates. All statistical analyses were performed using R v4.3.1 (www.r-project.org). Statistical significance was determined using Student’s *t*- tests: paired *t*-tests were used for pairwise comparisons between two different time points within a time-series dataset, and unpaired *t*-tests for comparisons between two different treatments.

## Acknowledgements

This study was supported by the National Research Foundation of Korea (NRF-2021R1A6A3A13044038, NRF-2021R1C1C1008303, NRF-2022R1A4A5031447, and RS-2024-00341771). The work at Monash University was supported by an NHMRC EL2 Fellowship (APP1178715) and an ARC Discovery Grant (DP210101595).

## FOOTNOTES

### Data, Materials, and Software Availability

The completed genomes and associated data, along with the raw transcriptome data, were deposited in NCBI’s GenBank and Sequence Read Archive (SRA) databases, respectively (Accession numbers: CP159910.1 and CP159800.1 for complete genomes, SRX25285529/SRX25285530 and SRX25286366/SRX25286367 for transcriptomes).

### Author contributions

HH and SY designed research; HH performed experiments; HH, TND, MJ, CG, and SY analyzed data; HH and TND prepared visualizations; and HH, TND, MJ, CG, and SY wrote the paper.

