## Supporting Information for "Hydrogen-dependent dissimilatory nitrate reduction to ammonium enables growth of *Campylobacterota* isolates"

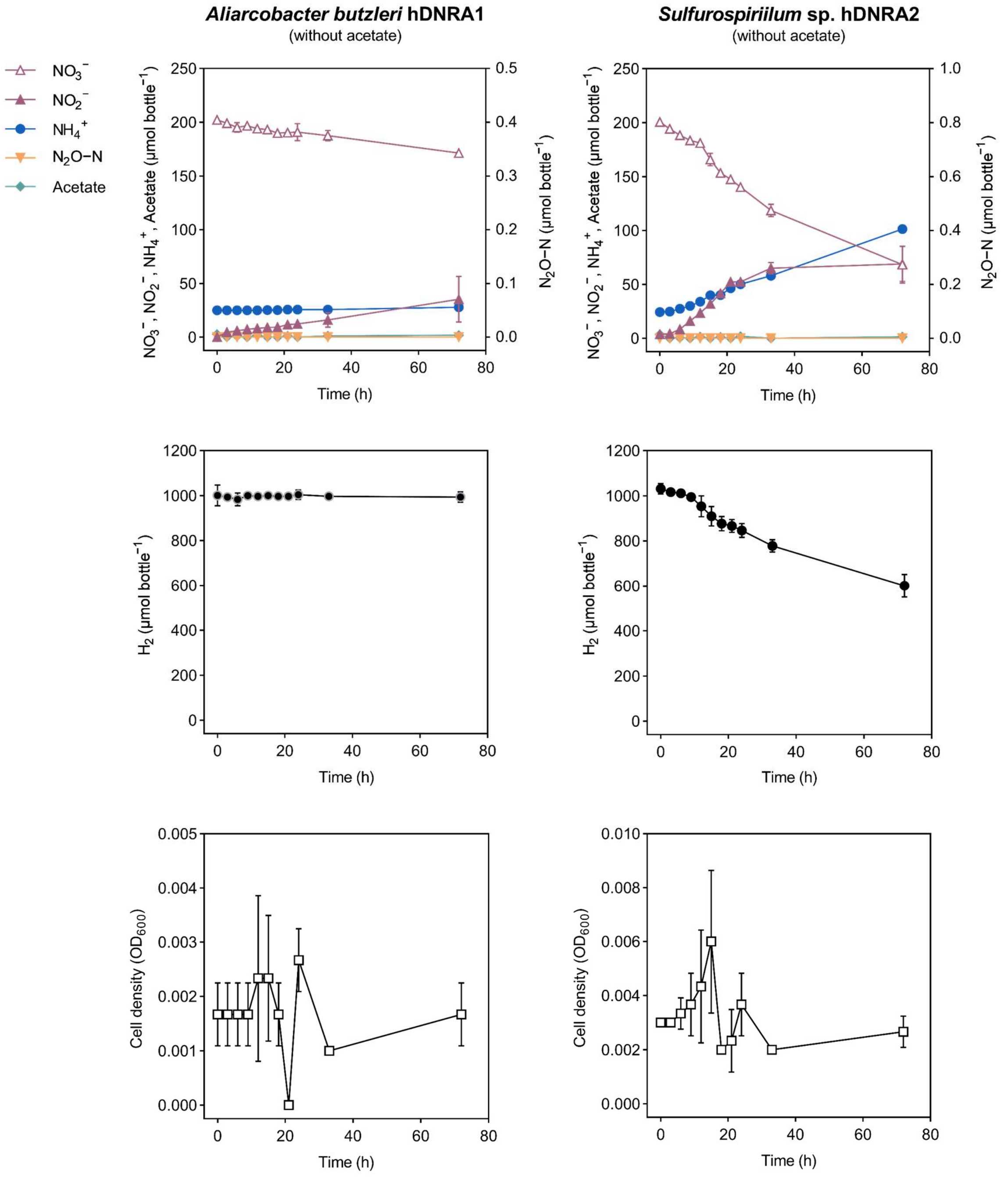


Fig. S1. Hydrogenotrophic DNRA activity in batch cultures of *A. butzleri* hDNRA1 (left column) and *Sulfurospirillum* sp. hDNRA2 (right column) incubated in the absence of organic carbon. Cultures were prepared with washed cells resuspended in fresh medium initially containing 2 mM $NO_3^-$ but lacking acetate, with an initial cell density of $OD_{600}$~0.002. The initial headspace contained 5% $H_2$ and 5% $CO_2$ (v/v). The concentrations of $NO_3^-$, $NO_2^-$, $NH_4^+$, $N_2O$-N, and $H_2$ in the bottle, along with cell density, were monitored over a 72-hour period. Each data point represents the mean of three biological replicates (*n*=3), with error bars indicating the standard deviations.

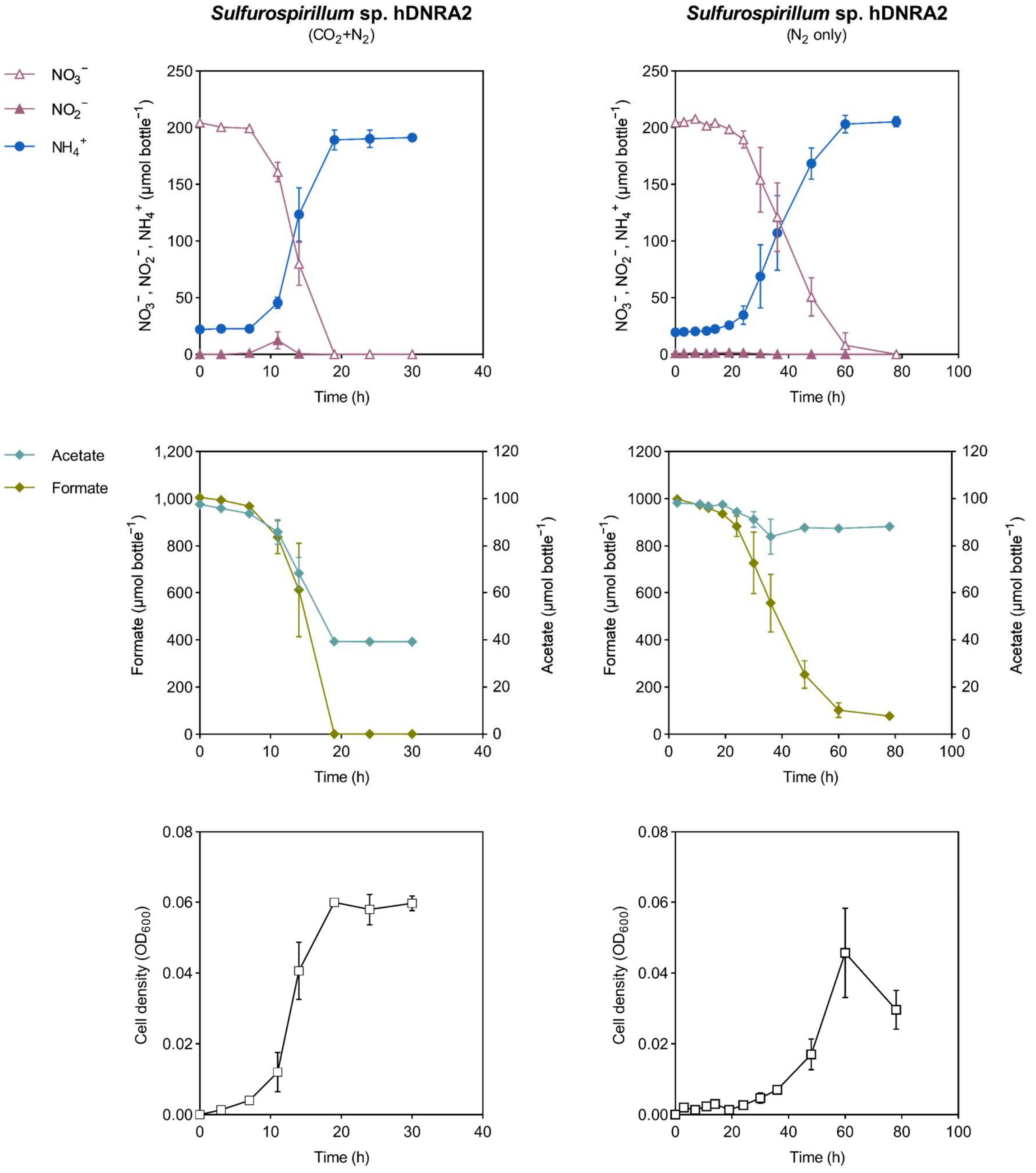


Fig. S2. Growth and $NO_3^-$/ $NO_2^-$ reduction by *Sulfurospirillum* sp. hDNRA2 observed in batch cultures initially amended with 2 mM $NO_3^-$, 10 mM formate, and 1 mM acetate, but without $H_2$ in the headspace. Cultures equilibrated with anoxic gases with (left column) and without (right column) 5% $CO_2$ (v/v) were examined. The concentrations of $NO_3^-$, $NO_2^-$, $NH_4^+$, acetate, formate, and cell density (measured as $OD_{600}$) were monitored until $NO_3^-$ and $NO_2^-$ were depleted. Each data point represents the mean of three biological replicates (*n*=3) with error bars indicating the standard deviations.

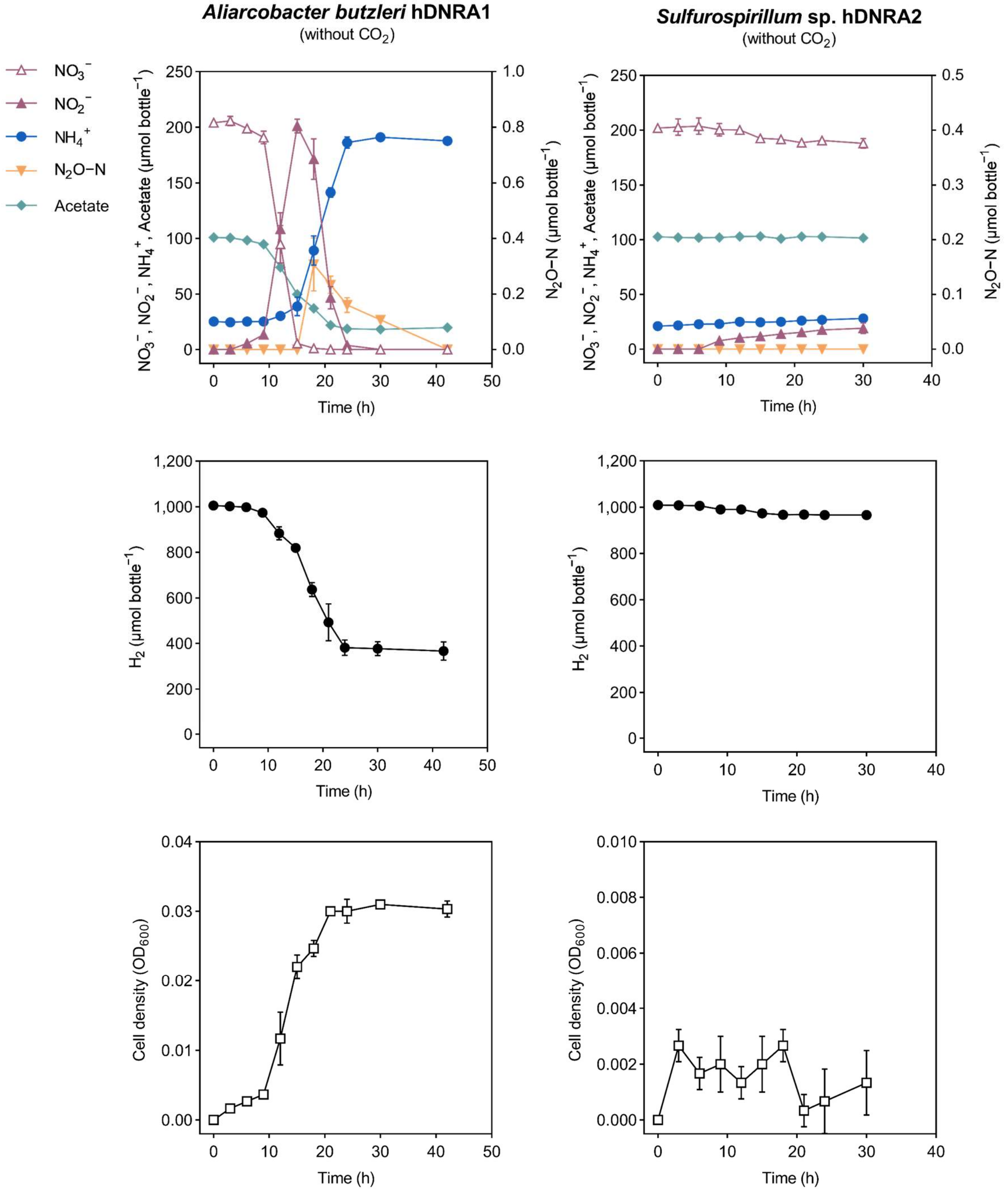


Fig. S3. Hydrogenotrophic DNRA activity in batch cultures of *A. butzleri* hDNRA1 (left column) and *Sulfurospirillum* sp. hDNRA2 (right column) incubated under initial absence of inorganic carbon. Transformation of $NO_3^-$ and associated $H_2$ consumption was examined in batch cultures prepared with 2 mM $NO_3^-$, 1 mM acetate, and a $CO_2$-free headspace (5% $H_2$/ 95% $N_2$; v/v). The concentrations of $NO_3^-$, $NO_2^-$, $NH_4^+$, $N_2O$-N, and $H_2$ in the bottle, as well as cell density, were monitored until the depletion of $NO_3^-$ and $NO_2^-$. Each data point represents the mean of three biological replicates (*n*=3), with error bars indicating the standard deviations.

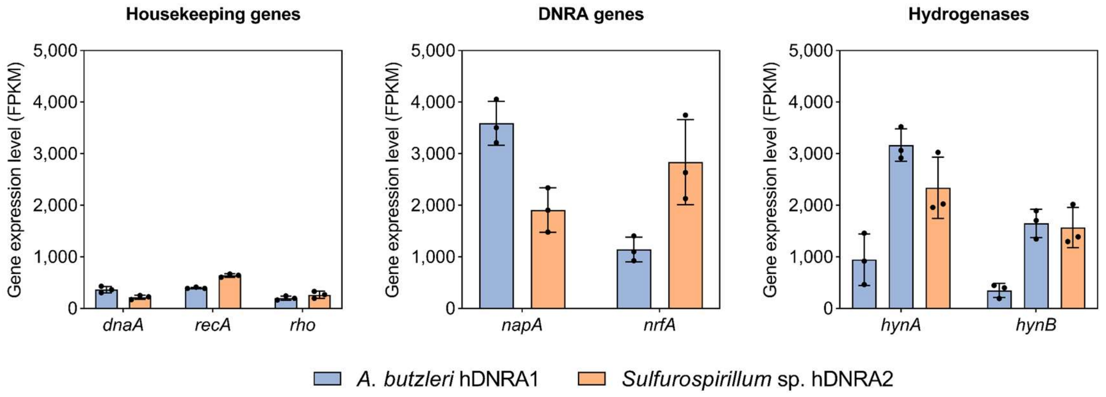


Fig. S4. Expression levels of the genes putatively involved in DNRA and $H_2$ oxidation compared with those of three single-copy housekeeping genes *dnaA*, *recA*, and *rho*, as computed from the sequenced transcriptomes of *A. butzleri* hDNRA1 and *Sulfurospirillum* sp. hDNRA2 grown on hydrogenotrophic DNRA. The expression levels of the genes encoding the periplasmic nitrate reductase (*napA*), cytochrome $c_{552}$ nitrite reductase (*nrfA*), and group 2d [NiFe]-hydrogenase (*hynAB*) are presented, along with those of the three single-copy housekeeping genes. Note that the complete genome of *A. butzleri* hDNRA1 contains two sets of *hynAB* genes, while all other genes examined here are single-copy in their respective genomes. Each bar length represents the mean of three biological replicates (*n*=3; each shown as a black dot) with error bars indicating the standard deviations.

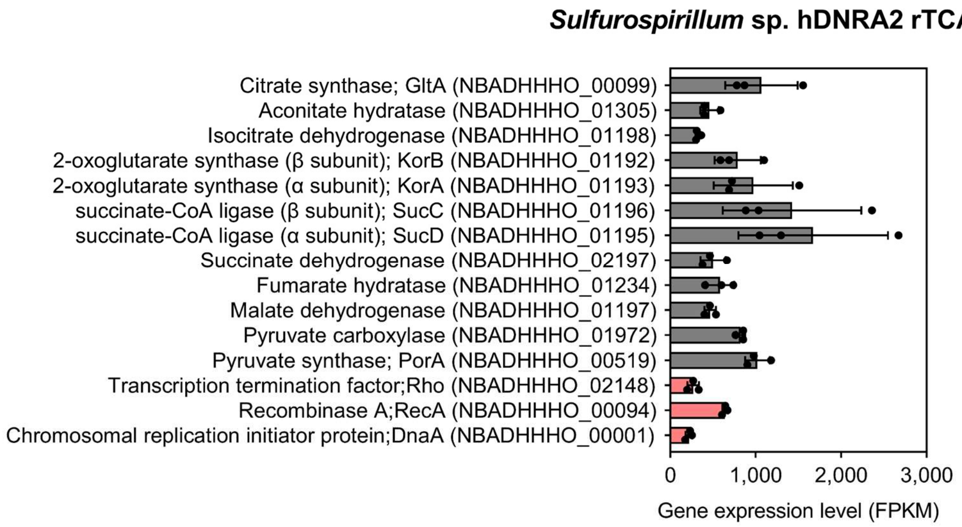


Fig. S5. Expression levels of the genes putatively involved in the reductive TCA (rTCA) pathway in transcriptome of *Sulfurospirillum* sp. hDNRA2 grown under hydrogenotrophic DNRA growth condition. Expression levels of single-copy housekeeping genes, *dnaA*, *recA*, and *rho* are also shown for comparison (red bar plots). Each bar length represents the mean of three biological replicates ($n$=3; each shown as a black dot) with error bars indicating the standard deviations.

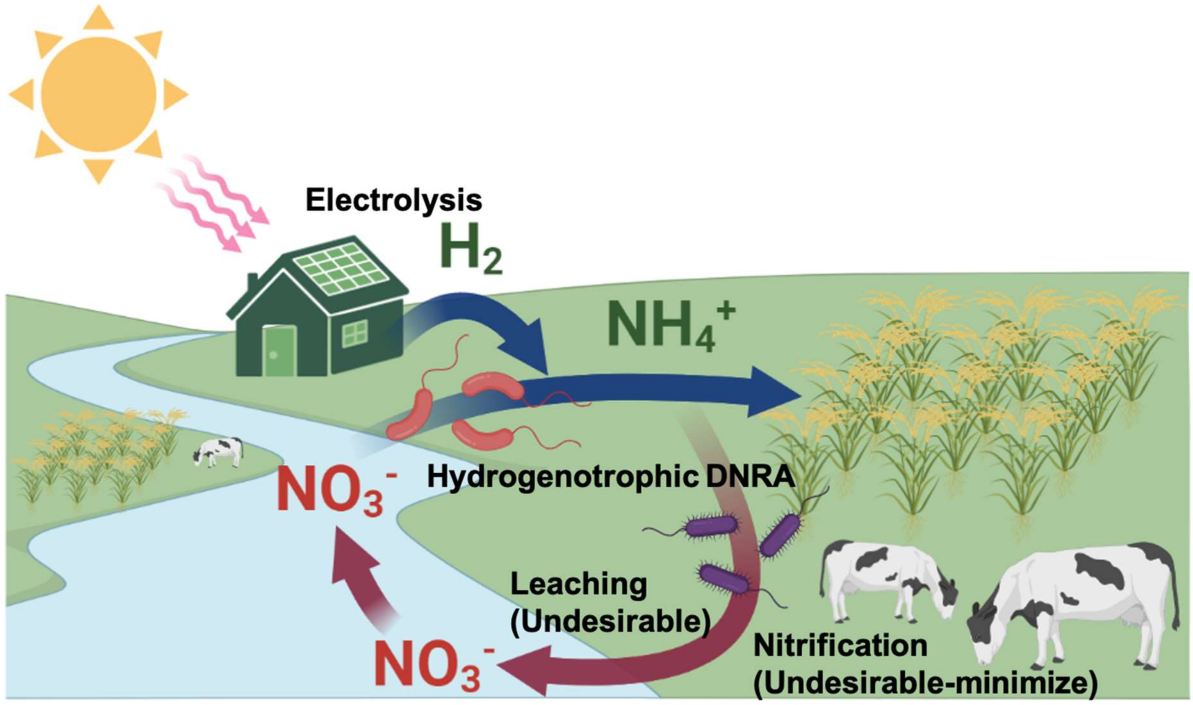


Fig. S6. Schematic of a potential biotechnological application of hydrogenotrophic DNRA for simultaneous $NO_3^-$ removal from freshwater bodies and $NH_4^+$ provision to agricultural soils as a green alternative to chemical fertilization.

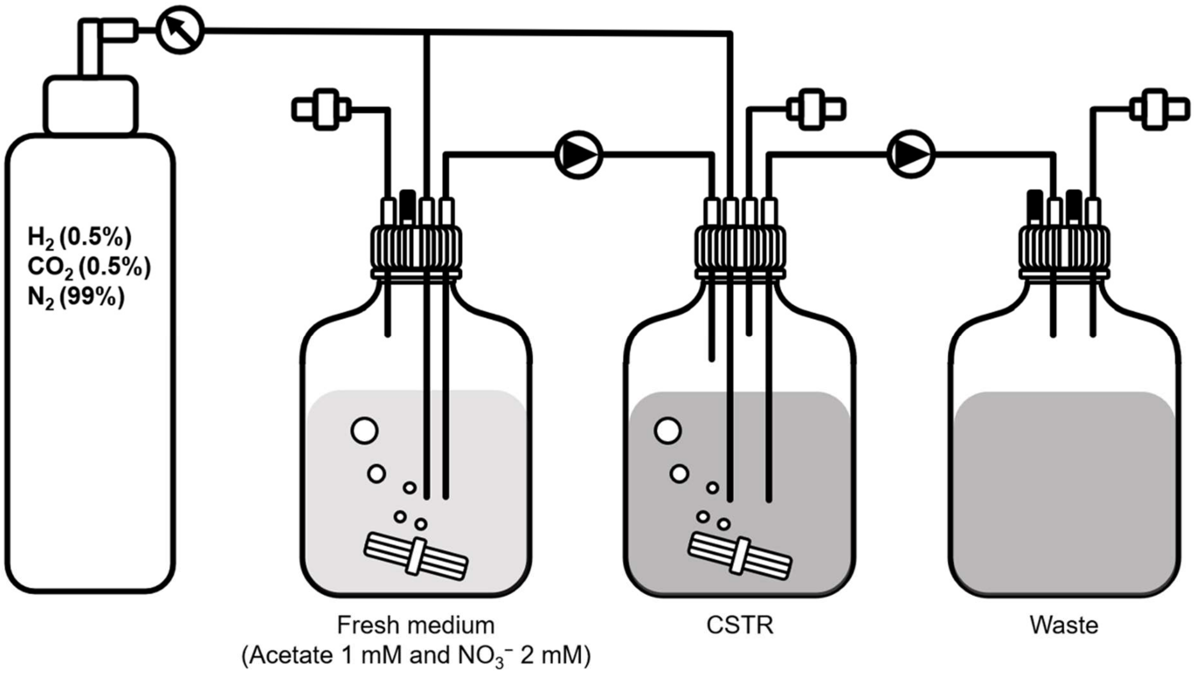


Fig. S7. Schematic depiction of the chemostat reactor used for the continuous cultivation of *A. butzleri* hDNRA1 and *Sulfurospirillum* sp. hDNRA2 under hydrogenotrophic DNRA conditions.

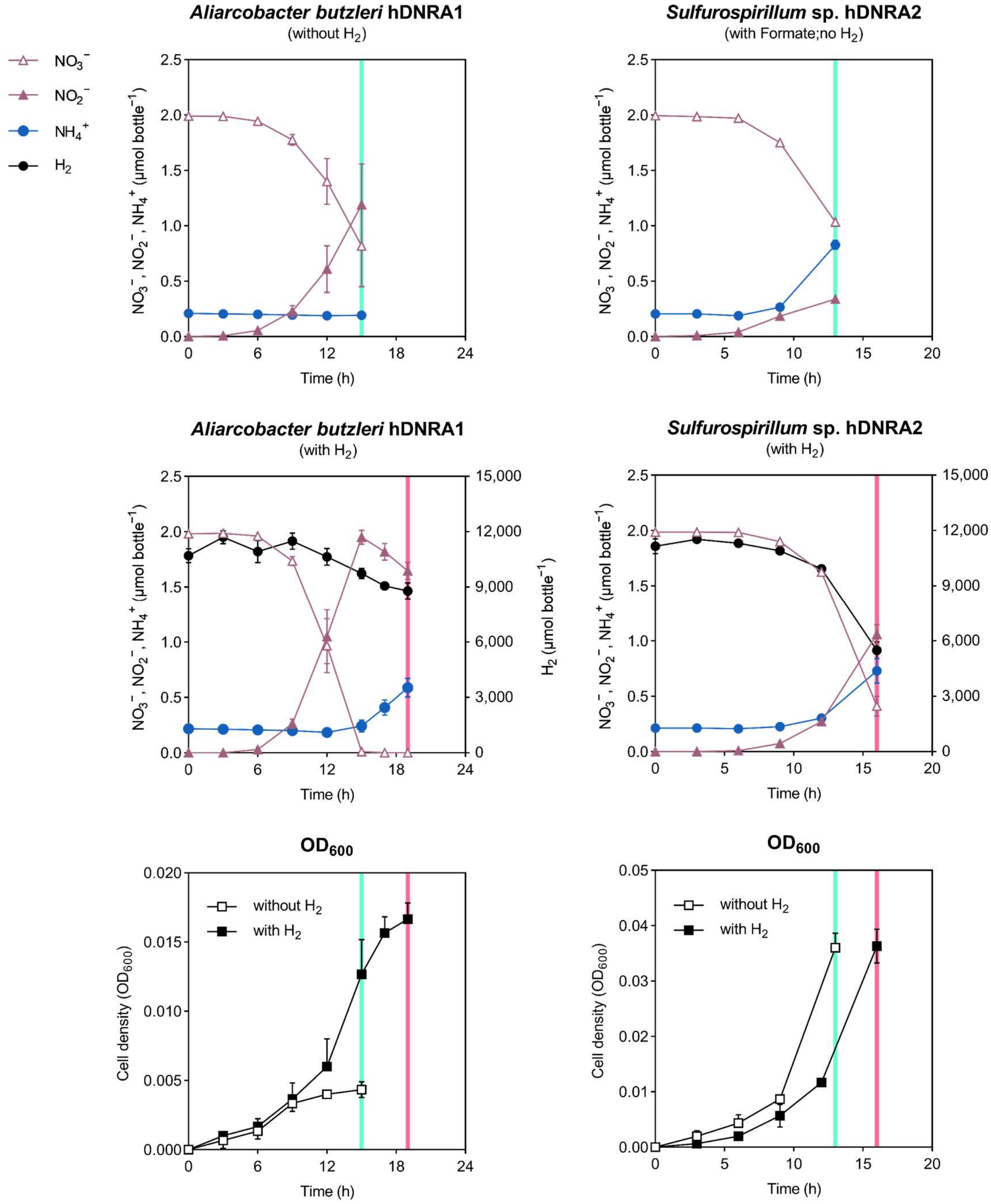


Fig. S8. Culture incubation and sample collection for transcriptome analysis. Batch-cultures of *A. butzleri* (left column) hDNRA1 and *Sulfurospirillum* sp. hDNRA2 (right column) were incubated in medium amended with 1 mM acetate and 2 mM $NO_3^-$. For the $H_2$-free *Sulfurospirillum* sp. hDNRA2 control, 10 mM formate was added, and cultures were equilibrated with either 5% $H_2$/5% $CO_2$/90% $N_2$ or 5% $CO_2$/95% $N_2$ (controls without $H_2$) mixed gas. The concentrations of $NO_3^-$, $NO_2^-$, $NH_4^+$, and $H_2$, along with cell density, were monitored until (i) active $NH_4^+$ production, (ii) $NO_3^-$ reduction and/or (iii) exponential growth was observed. Vertical lines indicate the time points for RNA sampling. Each data point represents the mean of three biological replicates (*n*=3), with samples separately collected, treated, and sequenced, and error bars indicating the standard deviations.

Table S1. Genome statistics

| **Organism** | *Aliarcobacter butzleri* hDNRA1 | *Sulfurospirillum* sp. hDNRA2 |
|---|---|---|
| **Genome size (bp)** | 2,306,496 | 2,684,720 |
| **GC percentage (%)** | 27.03 | 43.96 |
| **CheckM completeness (%)** | 100.0 | 100.0 |
| **CheckM contamination (%)** | 0.41 | 0.46 |
| **Number of contigs** | 1 | 2 (including 1 plasmid) |
| **Number of genes predicted** | 2,329 | 2,672 |
| **Number of protein coding genes** | 2,263 | 2,619 |
| **Number of tRNA genes** | 56 | 47 |
| **Number of rRNA genes** | 10 | 6 |

Table S2. List of functional genes relevant to nitrogen and hydrogen metabolisms in *Aliarcobacter butzleri* hDNRA1 and *Sulfurospirillum* sp. hDNRA2.

| **Gene name** | **Locus tag** | **Gene product**[a] | **Accession** | **E-value** | **EC number** | **Length (bp)** |
|---|---|---|---|---|---|---|
| *Aliarcobacter butzleri* hDNRA1 | | | | | | |
| Nitrogen metabolism | | | | | | |
| $NO_3^-$ reduction to $NO_2^-$ | | | | | | |
| *napD* | GODOFOPP_00358 | Chaperone NapD | COG3062 | 1.52e-21 | - | 384 |
| *napL* | GODOFOPP_00359 | WD40 repeat, putative periplasmic protein | COG2319 | 1.13e-08 | - | 960 |
| *napF* | GODOFOPP_00360 | Ferredoxin-type protein NapF | PRK10194 | 5.19e-21 | - | 477 |
| *napB* | GODOFOPP_00361 | Periplasmic nitrate reductase, small subunit (cytochrome *c*-type subunit) | COG3043 | 7.68e-40 | - | 606 |
| *napH* | GODOFOPP_00362 | Ferredoxin-type protein NapH | TIGR02163 | 4.26e-107 | - | 807 |
| *napG* | GODOFOPP_00363 | Ferredoxin-type protein NapG | TIGR00397 | 7.97e-68 | - | 813 |
| *napA* | GODOFOPP_00364 | Periplasmic nitrate reductase, larges subunit | TIGR01706 | 0.0 | 1.9.6.1 | 2,811 |
| $NO_2^-$ reduction to $NH_4^+$ | | | | | | |
| *nrfA* | GODOFOPP_00353 | Cytochrome *c* nitrite reductase (ammonia forming), large subunit | PRK11125 | 0.0 | 1.7.2.2 | 1,491 |
| *nrfH* | GODOFOPP_00354 | Cytochrome *c* nitrite reductase (ammonia forming), small subunit | TIGR03153 | 1.79e-52 | - | 531 |
| $N_2O$ reduction to $N_2$ | | | | | | |
| *nosZ* | GODOFOPP_00989 | Nitrous oxide reductase | COG4263 | 0.0 | 1.7.2.4 | 2,592 |
| *HP* | GODOFOPP_00990 | Hypothetical protein | - | - | - | 765 |
| *nosD* | GODOFOPP_00991 | Nitrous oxide reductase family maturation protein NosD | TIGR04247 | 1.54e-144 | - | 1,227 |
| - | GODOFOPP_00992 | Ferredoxin-type protein NapG | PRK09476 | 2.23e-48 | - | 717 |
| - | GODOFOPP_00993 | Cytochrome $c_{553}$ | COG2863 | 4.1e-10 | - | 558 |
| - | GODOFOPP_00994 | Cytochrome $c_{553}$ | COG2863 | 2.45e-13 | - | 444 |
| - | GODOFOPP_00995 | Ferredoxin-type protein NapH | PRK09477 | 5.84e-98 | - | 894 |
| - | GODOFOPP_00996 | Putative ABC transporter ATP-binding protein | TIGR04521 | 4.11e-34 | - | 639 |
| *nosL* | GODOFOPP_00997 | Nitrous oxide reductase accessory protein NosL | COG4314 | 3.51e-21 | - | 447 |
| *nosY* | GODOFOPP_00998 | ABC-type transport system involved in multi-copper enzyme maturation, permease component | COG1277 | 2.28e-14 | - | 828 |
| Hydrogenases[b] | | | | | | |
| *huaS* | GODOFOPP_01010 | Group 2d [NiFe]-hydrogenase, small subunit | COG1740 | 3.52e-61 | - | 894 |
| *huaL* | GODOFOPP_01011 | Group 2d [NiFe]-hydrogenase, large subunit | COG0374 | 2.49e-71 | - | 1,350 |
| *hynA* | GODOFOPP_01012 | Group 1b [NiFe]-hydrogenase, small subunit | COG1740 | 0.0 | - | 1,236 |
| *hynB* | GODOFOPP_01013 | Group 1b [NiFe]-hydrogenase, large subunit | COG0374 | 0.0 | - | 1,743 |
| *hynC* | GODOFOPP_01014 | [NiFe]-hydrogenase cytochrome *b* subunit | COG1969 | 2.14e-64 | - | 714 |

| | | | | | | |
|---|---|---|---|---|---|---|
| *hupD* | GODOFOPP_01015 | [NiFe]-hydrogenase maturation factor | COG0680 | 2.10e-32 | - | 591 |
| *hypF* | GODOFOPP_01016 | [NiFe]-hydrogenase maturation factor HypF (carbamoyltransferase) | COG0068 | 1.64e-07 | - | 1,533 |
| *hynA* | GODOFOPP_01019 | Group 1b [NiFe]-hydrogenase, small subunit | COG1740 | 0.0 | - | 1,179 |
| *hynB* | GODOFOPP_01020 | Group 1b [NiFe]-hydrogenase, large subunit | COG0374 | 0.0 | - | 1,731 |
| *hynC* | GODOFOPP_01021 | [NiFe]-hydrogenase cytochrome *b* subunit | COG1969 | 7.57e-41 | - | 666 |
| *hyaD* | GODOFOPP_01022 | [NiFe]-hydrogenase maturation factor | COG0680 | 7.30e-30 | - | 579 |
| *HP* | GODOFOPP_01023 | Hypothetical protein | - | - | - | 1,620 |
| *hypF* | GODOFOPP_01024 | [NiFe]-hydrogenase maturation factor HypF (carbamoyltransferase) | COG0068 | 0.0 | - | 2,247 |
| *hypB* | GODOFOPP_01048 | Hydrogenase/urease maturation factor HypB, $Ni^{2+}$-binding GTPase | COG0378 | 1.55e-97 | - | 816 |
| *hypC* | GODOFOPP_01049 | Hydrogenase maturation factor HybG, HypC/HupF family | COG0298 | 1.40e-29 | - | 282 |
| *hypD* | GODOFOPP_01050 | Hydrogenase maturation factor HypD | COG0409 | 0.0 | - | 1,125 |
| *hypE* | GODOFOPP_01055 | Carbamoyl dehydratase HypE (hydrogenase maturation factor) | COG0309 | 3.71e-162 | - | 999 |
| *hypA* | GODOFOPP_01056 | Hydrogenase maturation factor HypA/HybF, metallochaperone involved in Ni insertion | COG0375 | 2.77e-39 | - | 342 |
| *Sulfurospirillum* sp. hDNRA2 | | | | | | |
| Nitrogen metabolism | | | | | | |
| $NO_3^-$ reduction to $NO_2^-$ | | | | | | |
| *napD* | NBADHHHO_01834 | Chaperone NapD | COG3062 | 2.32e-23 | - | 354 |
| *napL* | NBADHHHO_01835 | WD40 repeat, putative periplasmic protein | COG2319 | 4.86e-18 | - | 948 |
| *napF* | NBADHHHO_01836 | Ferredoxin-type protein NapF | PRK10194 | 4.16e-24 | - | 504 |
| *napB* | NBADHHHO_01837 | Periplasmic nitrate reductase, small subunit (cytochrome *c*-type subunit) | COG3043 | 5.41e-54 | - | 525 |
| *napH* | NBADHHHO_01838 | Ferredoxin-type protein NapH | TIGR02163 | 2.29e-113 | - | 825 |
| *napG* | NBADHHHO_01839 | Ferredoxin-type protein NapG | TIGR00397 | 1.96e-62 | - | 822 |
| *napA* | NBADHHHO_01840 | Periplasmic nitrate reductase catalytic/large subunit | TIGR01706 | 0.0 | 1.9.6.1 | 2,790 |
| $NO_2^-$ reduction to $NH_4^+$ | | | | | | |
| *nrfA* | NBADHHHO_01869 | Cytochrome *c* nitrite reductase (ammonia forming), large subunit | PRK11125 | 0.0 | 1.7.2.2 | 1,548 |
| *nrfH* | NBADHHHO_01870 | Cytochrome *c* nitrite reductase (ammonia forming), small subunit | TIGR03153 | 4.11e-62 | - | 534 |
| $N_2O$ reduction to $N_2$ | | | | | | |
| *nosZ* | NBADHHHO_01560 | Nitrous-oxide reductase | COG4263 | 0.0 | 1.7.2.4 | 2,595 |
| *nosY* | NBADHHHO_01747 | ABC-type transport system involved in multi-copper enzyme maturation, permease component | COG1277 | 5.57e-15 | - | 828 |
| *nosL* | NBADHHHO_01748 | Nitrous oxide reductase accessory protein NosL | COG4314 | 3.78e-11 | - | 504 |

| | | | | | | |
|---|---|---|---|---|---|---|
| *HP* | NBADHHHO_01749 | Hypothetical protein | - | - | - | 966 |
| *lolD* | NBADHHHO_01750 | ABC-type lipoprotein export system, ATPase component | COG1136 | 1.10e-95 | - | 669 |
| *lolC* | NBADHHHO_01751 | ABC-type transport system involved in lipoprotein release, permease component | COG4591 | 2.45e-27 | - | 1,194 |
| *HP* | NBADHHHO_01752 | Hypothetical protein | - | - | - | 318 |
| *nosL* | NBADHHHO_01753 | Nitrous oxide reductase accessory protein NosL | COG4314 | 2.62e-22 | - | 456 |
| - | NBADHHHO_01754 | Putative ABC transporter ATP-binding protein | pfam00005 | 1.09e-31 | - | 639 |
| - | NBADHHHO_01755 | Ferredoxin-type protein NapH | TIGR02163 | 6.19e-91 | - | 906 |
| - | NBADHHHO_01756 | Cytochrome $c_{553}$ | COG2863 | 3.56e-16 | - | 495 |
| - | NBADHHHO_01757 | Cytochrome $c_{553}$ | COG2863 | 9.39e-13 | - | 579 |
| - | NBADHHHO_01758 | Ferredoxin-type protein NapG | TIGR00397 | 1.82e-32 | - | 726 |
| *nosD* | NBADHHHO_01759 | Nitrous oxide reductase family maturation protein NosD | TIGR04247 | 1.38e-151 | - | 1,308 |
| *HP* | NBADHHHO_01760 | Hypothetical protein | - | - | - | 795 |
| *nosZ* | NBADHHHO_01761 | Nitrous-oxide reductase | COG4263 | 0.0 | 1.7.2.4 | 2,601 |
| Hydrogenase | | | | | | |
| *hypA* | NBADHHHO_01203 | Hydrogenase maturation factor HypA/HybF, metallochaperone involved in Ni insertion | COG0375 | 4.10e-42 | - | 342 |
| *hypE* | NBADHHHO_01204 | Carbamoyl dehydratase HypE (hydrogenase maturation factor) | COG0309 | 1.62e-166 | - | 1,005 |
| *hypD* | NBADHHHO_01205 | Hydrogenase maturation factor HypD | COG0409 | 0.0 | - | 1,125 |
| *hypC* | NBADHHHO_01206 | Hydrogenase maturation factor HybG, HypC/HupF family | COG0298 | 2.00e-32 | - | 270 |
| *hypB* | NBADHHHO_01207 | Hydrogenase/urease maturation factor HypB, $Ni^{2+}$-binding GTPase | COG0378 | 1.21e-124 | - | 837 |
| *hypF* | NBADHHHO_01382 | [NiFe]-hydrogenase maturation factor HypF (carbamoyltransferase) | COG0068 | 0.0 | - | 2,220 |
| *HP* | NBADHHHO_01383 | Hypothetical protein | - | - | - | 1,638 |
| *hupD* | NBADHHHO_01384 | [NiFe]-hydrogenase maturation factor | COG0680 | 3.00e-32 | - | 540 |
| *hynC* | NBADHHHO_01385 | [NiFe]-hydrogenase cytochrome *b* subunit | COG1969 | 7.71e-38 | - | 678 |
| *hynB* | NBADHHHO_01386 | Group 1b [NiFe]-hydrogenase, large subunit | COG0374 | 0.0 | - | 1,749 |
| *hynA* | NBADHHHO_01387 | Group 1b [NiFe]-hydrogenase, small subunit | COG1740 | 0.0 | - | 1,158 |
| *huaL* | NBADHHHO_01388 | Group 2d [NiFe]-hydrogenase, large subunit | COG0374 | 2.05e-71 | - | 1,344 |
| *huaS* | NBADHHHO_01389 | Group 2d [NiFe]-hydrogenase, small subunit | COG1740 | 5.81e-57 | - | 897 |
| *hycG* | NBADHHHO_01479 | Group 4c [NiFe]-hydrogenase, small subunit | COG3260 | 4.87e-51 | - | 411 |
| - | NBADHHHO_01480 | Formate hydrogenlyase complex iron-sulfur subunit | PRK12387 | 7.08e-16 | - | 426 |
| *hycE1* | NBADHHHO_01481 | NADH-quinone oxidoreductase subunit C | COG3262 | 2.05e-17 | - | 495 |
| *hycE2* | NBADHHHO_01482 | Group 4c [NiFe]-hydrogenase, large subunit | COG3261 | 1.31e-161 | - | 1,089 |
| *hycB* | NBADHHHO_01483 | Fe-S-cluster-containing hydrogenase component 2 | cd10554 | 2.41e-55 | - | 474 |
| *hypA* | NBADHHHO_01484 | Hydrogenase maturation factor HypA/HybF, metallochaperone | COG0375 | 5.53e-38 | - | 363 |

| | | | | | | |
|---|---|---|---|---|---|---|
| | | involved in Ni insertion | | | | |
| *gltD* | NBADHHHO_01485 | Glutamate synthase (NADPH), beta chain | COG0493 | 0.0 | 1.4.1.13 | 1,425 |
| - | NBADHHHO_01844 | Putative Fe-S cluster-containing hydrogenase component | COG1142 | 6.55e-44 | - | 654 |
| *hyfB* | NBADHHHO_01845 | Putative multi-subunit $Na^+/H^+$ antiporter | COG0651 | 6.05e-86 | - | 1,941 |
| *hyfC* | NBADHHHO_01846 | Formate hydrogenlyase subunit HyfC | COG0650 | 3.97e-80 | - | 927 |
| *hyfE* | NBADHHHO_01847 | Hydrogenase membrane subunit HyfE | COG4237 | 2.73e-48 | - | 654 |
| *hyfB* | NBADHHHO_01848 | Putative multi-subunit $Na^+/H^+$ antiporter | COG0651 | 2.05e-93 | - | 1,470 |
| *hyfE* | NBADHHHO_01849 | Group 4a [NiFe]-hydrogenase large subunit | COG3261 | 0.0 | - | 1,740 |
| - | NBADHHHO_01850 | Formate hydrogenlyase complex iron-sulfur subunit | PRK12387 | 5.83e-98 | - | 540 |
| *hyfG* | NBADHHHO_01851 | Group 4a [NiFe]-hydrogenase small subunit | COG3260 | 1.90e-88 | - | 822 |
| *hyfH* | NBADHHHO_01852 | Formate hydrogenlyase maturation protein | PRK15084 | 5.97e-09 | - | 360 |
| *hyfD* | NBADHHHO_01853 | Endopeptidases belonging to membrane-bound hydrogen evolving hydrogenase group | cd06067 | 1.99e-45 | - | 447 |

[a]Gene annotation was performed using NCBI's non-redundant sequence database as the reference database
[b]Hydrogenase classification was in accordance with HydDB (https://services.birc.au.dk/hyddb/)

Table S3. List of genes in the aspartate ammonia-lyase cluster

| **Gene name** | **Locus tag** | **Gene product**[a] | **Accession** | **E-value** | **EC number** | **Length (bp)** |
|---|---|---|---|---|---|---|
| *hydF* | NBADHHHO_01802 | [FeFe]-hydrogenase H-cluster maturation GTPase HydF | TIGR03918 | 0.0 | - | 1,227 |
| *hydE* | NBADHHHO_01803 | [FeFe]-hydrogenase H-cluster radical SAM maturase HydE | TIGR03956 | 1.87e-127 | - | 1,068 |
| *aspA* | NBADHHHO_01804 | Aspartate ammonia-lyase | COG1027 | 0.0 | 4.3.1.1 | 1,392 |
| *hydG* | NBADHHHO_01805 | [FeFe]-hydrogenase H-cluster radical SAM maturase HydG | TIGR03955 | 2.03e-156 | - | 1473 |
| - | NBADHHHO_01806 | Cytochrome *b* subunit | COG2864 | 1.10e-22 | - | 690 |
| *hydB* | NBADHHHO_01807 | Group A [FeFe]-hydrogenase, small subunit | pfam02256 | 2.41e-11 | - | 363 |
| *hydA* | NBADHHHO_01808 | Group A [FeFe]-hydrogenase, large subunit | TIGR02512 | 8.78e-138 | - | 1,353 |

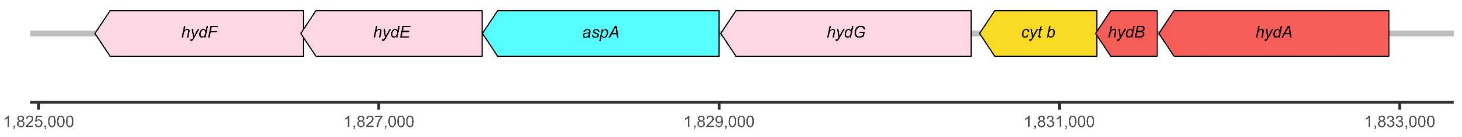


[a]Gene annotation was performed using NCBI's non-redundant sequence database as the reference database
